# Dynamics of bacterial cell division: Z ring condensation is essential for cytokinesis

**DOI:** 10.1101/2020.06.30.180737

**Authors:** Georgia R. Squyres, Matthew J. Holmes, Sarah R. Barger, Betheney R. Pennycook, Joel Ryan, Victoria T. Yan, Ethan C. Garner

## Abstract

How proteins in the bacterial cell division complex (the divisome) coordinate to divide bacteria remains unknown. To explore how these proteins collectively function, we conducted a complete dynamic characterization of the proteins involved, and then examined the function of FtsZ binding proteins (ZBPs) and their role in cytokinesis. We find that the divisome consists of two dynamically distinct subcomplexes: stationary ZBPs that transiently bind to treadmilling FtsZ filaments, and a directionally-moving complex that includes cell wall synthases. FtsZ filaments treadmill at steady state and the ZBPs have no effect on filament dynamics. Rather, ZBPs bundle FtsZ filaments, condensing them into Z rings. Z ring condensation increases the recruitment of cell wall synthesis enzymes to the division site, and this condensation is necessary for cytokinesis.

## Main Text

The mechanism by which bacteria divide remains poorly understood. In most bacteria, division begins when filaments of FtsZ, a tubulin homolog, form a “Z ring” at midcell (*1*). The Z ring then recruits other cell division proteins, collectively called the divisome (Fig 1A). The first group of these proteins (early proteins) arrives concurrently with FtsZ and includes the actin homolog FtsA and several other FtsZ binding proteins (ZBPs): EzrA, SepF, and ZapA. The second group of integral membrane proteins (late proteins) is then recruited, including DivIB, DivIC, and FtsL, and the cell wall synthesis enzymes Pbp2B and FtsW (*2, 3*). During cytokinesis, the Z ring constricts while the associated cell wall synthesis enzymes build a septum that divides the cell in half (*4*). Recent work has shown that FtsZ filaments treadmill around the division plane, moving at the same rate as the transpeptidase Pbp2B. (*5, 6*) (Movie S1). FtsZ treadmilling dynamics are critical for cell division: In *Bacillus subtilis*, the rate of treadmilling limits Pbp2B motion, the rate of septal cell wall synthesis, and the overall rate of septation (*5*).

**Fig. 1:**
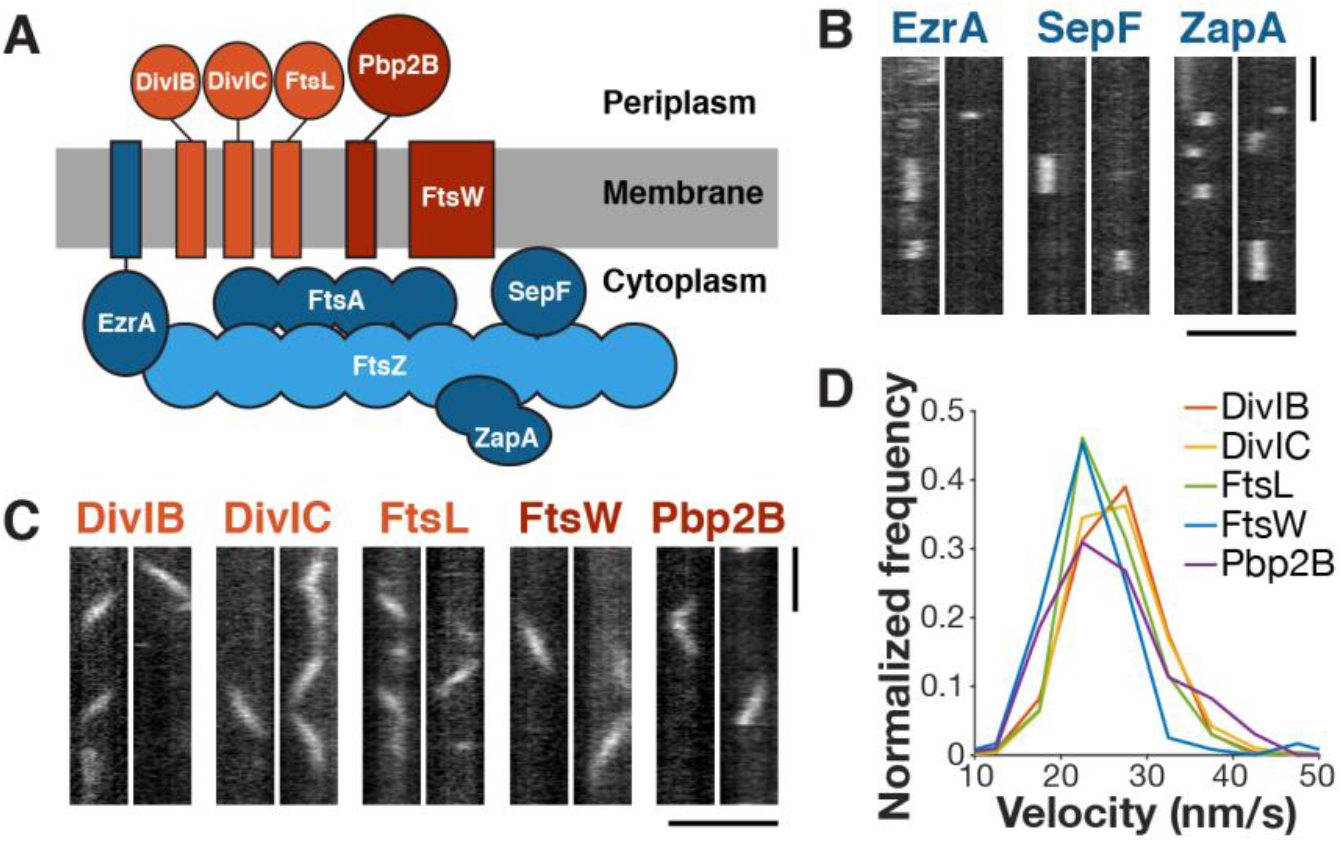
The divisome consists of two dynamic subcomplexes. **A** Schematic of divisome proteins in *B. subtilis*, with early-arriving proteins in red and late-arriving proteins in blue. *Light blue*: FtsZ filament, *dark blue*: FtsZ binding proteins, *light red*: trimeric complex, *dark red*: cell wall synthesis enzymes. All blue proteins are stationary, and all red proteins move directionally at the same velocity. **B** Kymographs of single molecules of stationary ZBPs at division sites. Each protein was tagged with a HaloTag and labeled with JF549-HTL for single-molecule imaging. Molecules were selected that colocalize with Z rings, visualized with epifluorescence imaging of mNeonGreen-FtsZ. Strains: EzrA: bMH42, SepF: bMH372, ZapA: bMH49. **C** Kymographs of single molecules of directionally-moving proteins at division sites. Each protein was tagged with a HaloTag and labeled with JF549-HTL for single-molecule imaging. Molecules are selected that colocalize with Z rings as above. Strains: DivIB: bAB366, DivIC: bAB367, FtsL: bGS65, FtsW: bAB368, Pbp2B: bGS31. **D** Velocity distributions of all directionally-moving proteins, measured from kymographs. Scale bars: horizontal: 2 μm, vertical: 1 min.

To understand how these proteins work to divide cells, we sought to build a dynamic characterization of how this multi-component machine functions in *B. subtilis*. We first worked to identify groups of divisome proteins that move together, then investigated how the FtsZ-associated proteins modulate FtsZ filaments, cell wall synthesis, and the overall process of cell division.

First, in order to understand which of the divisome proteins in *B. subtilis* associate with each other and work together, we characterized their dynamics using single-molecule imaging, as associated proteins should have similar motions. We expressed HaloTag fusions of each protein either as a sole copy or, in the case of SepF, at low levels from an ectopic site. Cells were sparsely labeled with JF549-HaloTag Ligand (*7*) and imaged with Total Internal Reflection Fluorescence Microscopy (TIRFM). Just as single molecules of FtsZ and FtsA are immobile (*5*), single molecules of the ZBPs were stationary (Fig 1B, Movie S2), consistent with their binding to stationary FtsZ subunits within treadmilling filaments. In contrast, the late proteins all moved directionally, with velocity distributions similar to Pbp2B (Fig 1C-D, Movie S3). The divisome-associated cell wall synthesis enzymes are known to function together (*8, 9*), and these data also show that the DivIB-DivIC-FtsL trimeric complex (*10*) remains persistently associated with these enzymes as they move around the division site. Thus, the divisome is composed of two distinct dynamic subcomplexes: 1) a directionally-moving group of periplasmic-facing membrane proteins that includes the cell wall synthesis enzymes, and 2) a group of cytoplasmic-facing proteins that bind to the stationary subunits within treadmilling FtsZ filaments.

Next, we investigated the function of the stationary subcomplex. We first examined FtsZ treadmilling, both to build a detailed characterization of its *in vivo* polymer dynamics and to provide a baseline for further investigation of how the ZBPs might affect its dynamics. In addition to visualizing FtsZ filament motion, we also measured FtsZ’s subunit lifetimes: While filament motion reports only treadmilling velocity, subunit lifetime measurements reflect both the treadmilling velocity and filament length (*11, 12*) (Fig 2A). Single-molecule imaging of FtsZ (Fig 2B, S1, Movie S4) gave a mean lifetime of 8.1 seconds (Fig 2C), corresponding to t½ = 5.6 s, similar to measurements of Z ring recovery after photobleaching (t½ = 8 ± 3 s) (*13*). Interestingly, FtsZ subunit lifetimes were higher at the division site than elsewhere (Fig 2D), possibly due to proteins such as Min and Noc that destabilize FtsZ filaments away from the division site (*14*). Confirming that our lifetime measurements report on FtsZ treadmilling, lifetimes increased with FtsZ(T111A), a mutant that treadmills more slowly, and decreased when treadmilling was accelerated by MciZ expression (*5*) (Fig 2E). We also characterized the dynamics of the rest of the stationary subcomplex and found that the lifetimes of FtsA and the ZBPs were similar to or slightly shorter than that of FtsZ (Fig 2F).

**Fig. 2:**
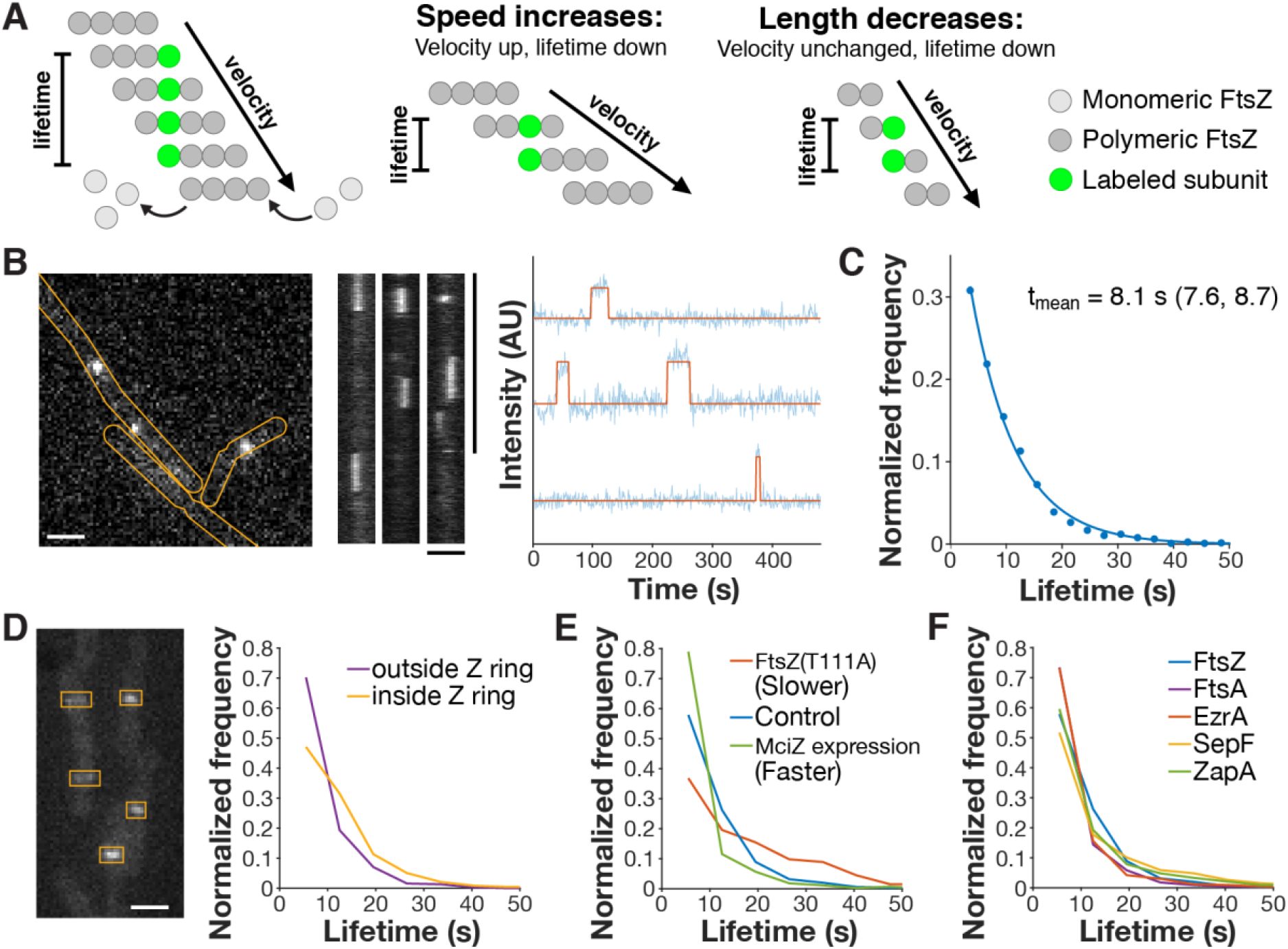
FtsZ lifetime reports treadmilling dynamics *in vivo*. **A** *Left*: Velocity and lifetime can be measured independently for a treadmilling filament. Velocity is measured by visualizing the motion of FtsZ filaments (dark gray circles), whereas lifetime is measured from the dwell time of a single labeled subunit (green circle) in the filament. *Center*: If the speed of a FtsZ filament is changed, this will result in a change in both the velocity and the lifetime that we measure. *Right*: If the length of a FtsZ filament is changed, the lifetime will change, but the velocity will not. Thus, lifetime reflects both treadmilling speed and filament length, whereas velocity is insensitive to filament length. **B** Lifetimes were measured by live-cell single-molecule TIRFM (*left*) of stationary FtsZ subunits (kymographs, *center*). FtsZ-HaloTag was induced with 20 μM IPTG for 2 hours and labeled with 40 pM JF549-HTL. Intensity traces were fit to a hidden Markov model to measure single-molecule lifetimes (*right*). Strain bAB309. **C** FtsZ subunit lifetime distribution, fit to a single exponential *f(t) = Ae^−τt^*. t_mean_: mean lifetime and 95% confidence interval, measured from this fit. Combined data from bAB309 and bGS104. **D** *Left*: mNeonGreen-Pbp2B was imaged by epifluorescence microscopy, and ROIs were drawn around Z rings. *Right*: Lifetime distributions of FtsZ inside and outside of Z rings, classified by colocalization with ROIs. Strain bGS104. **E** FtsZ subunit lifetime distributions in conditions that change treadmilling speed. Slower: GTP hydrolysis-deficient mutant FtsZ(T111A), Faster: MciZ expression. Strains: Slower: bGS109, Control: bAB309 and bGS104, Faster: bGS328. **F** Lifetime distributions of FtsZ, FtsA, and the ZBPs. HaloTag fusions of each protein were labeled with JF549-HTL for single-molecule imaging. Strains: FtsZ: bAB309 and bGS104, FtsA: bAB213, EzrA: bMH03, SepF: bMH332, ZapA: bMH28. Scale bars: horizontal: 2 μm, vertical: 1 min.

Combining our lifetime measurements with treadmilling velocity gives insight into the *in vivo* properties of FtsZ filaments. First, we estimate that the average FtsZ filament is ~230 nm long, similar to the lengths of purified FtsZ filaments measured by electron microscopy (200 ± 75 nm) (*15*). Second, assuming that treadmilling filaments elongate with a diffusion-limited on-rate of 5 μM-1 s-1 (*16, 17*), the concentration of free FtsZ monomer in cells can be estimated to be ~1.3 μM, similar to the critical concentration of FtsZ measured *in vitro* (1-1.5 μM) (*15*). This agreement between the critical concentration *in vitro* and *in vivo* suggests that the dynamics of FtsZ polymers *in vivo* are intrinsic to the polymer itself and unaffected by other factors. However, past work has suggested that FtsZ filament dynamics are modulated by other proteins *in vivo* (*2*).

To resolve this, we next investigated whether ZBPs affected FtsZ dynamics *in vivo*. ZBPs have been proposed to regulate FtsZ filament dynamics, stability, GTPase activity, and bundling, based both on *in vitro* biochemistry and *in vivo* genetic interactions (*18–27*). We measured FtsZ’s treadmilling velocity and subunit lifetime in strains as we changed the levels of individual ZBPs by deletion or overexpression (Fig 3A, S2-4, Table S3). None of these ZBP perturbations had any effect on FtsZ’s treadmilling velocity. Knockouts of ZapA or SepF also did not change FtsZ subunit lifetime, and overexpression had only subtle effects (Fig S3, Table S4). However, Δ*ezrA* cells had increased lifetimes, consistent with previous FRAP measurements of FtsZ-GFP (*13*), and EzrA overexpression decreased lifetime in a dose-dependent manner (Fig 3B, S4). EzrA’s effect of decreasing subunit lifetime without changing treadmilling speed suggest that it shortens FtsZ filaments, which we confirmed using Structured Illumination-TIRF (SIM-TIRF) (Fig 3C, Movie S5); however, it does so without changing FtsZ’s treadmilling dynamics, likely by binding to FtsZ monomers and sequestering them from the polymerizing pool (Supplementary Text 1). Thus, the ZBPs individually do not affect FtsZ treadmilling *in vivo*.

**Fig. 3:**
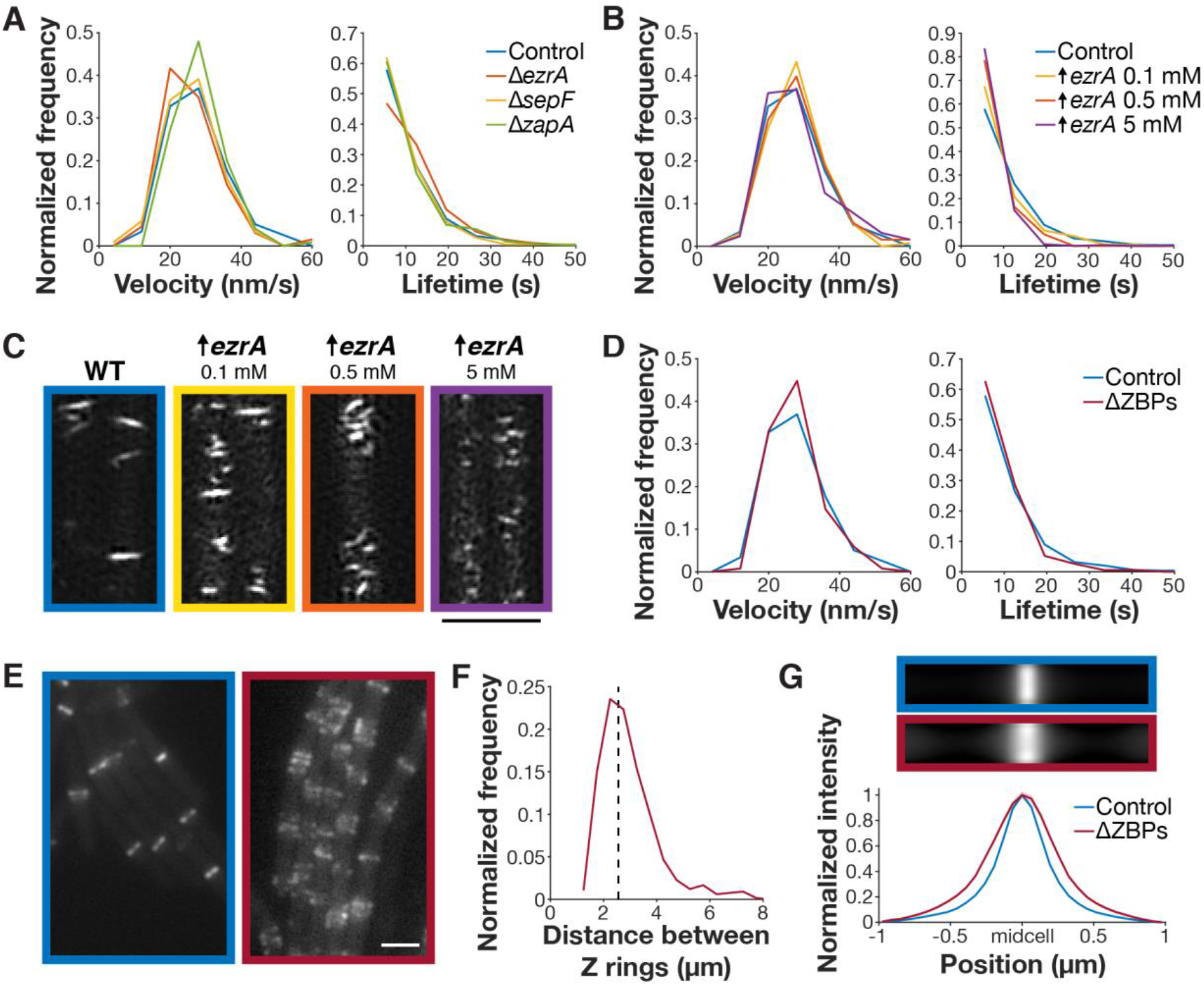
The ZBPs affect Z ring morphology but not dynamics. **A** *Left*: FtsZ filament velocity distributions in cells with individual ZBPs deleted. For velocity measurements, FtsZ-mNeonGreen was induced with 20 μM IPTG for 2 hours, imaged by TIRFM, and analyzed from kymographs. *Right*: FtsZ subunit lifetime distributions in cells with individual ZBPs knocked out. For lifetime measurements, FtsZ-HaloTag was induced with 20 μM IPTG for 2 hours and labeled with 40 pM JF549-HTL. Strains, *left*: control: bAB219, Δ*ezrA*: bGS256, Δ*sepF*: bGS254, Δ*zapA*: bGS250. Strains, *right*: control: bAB309 and bGS104, Δ*ezrA*: bGS167, Δ*sepF*: bGS304, Δ*zapA*: bGS141. **B** *Left*: FtsZ filament velocity distributions in cells with EzrA overexpressed. A second copy of EzrA was induced at the indicated xylose concentration for 2 hours. *Right*: FtsZ subunit lifetime distributions in cells with EzrA overexpressed. Strains, *left*: control: bAB219, *ezrA*↑: bGS263. Strains, *right*: control: bAB309 and bGS104, *ezrA*↑: bGS157. **C** FtsZ filaments in cells overexpressing EzrA, visualized by SIM-TIRF imaging. A second copy of EzrA was induced at the indicated xylose concentration for 2 hours. Strains: control: bAB219, *ezrA*↑: bGS263. **D** *Left*: FtsZ filament velocity distributions in cells with all ZBPs removed. ΔZBPs cells had *sepF* and *zapA* deleted and a xylose-inducible *ezrA*, which was depleted for 7 hours before imaging. *Right*: FtsZ subunit lifetime distributions in cells with all ZBPs removed. Strains, *left*: control: bAB219, ΔZBPs: bGS308. Strains, *right*, control: bAB309 and bGS104, ΔZBPs: bGS331. E Z rings in control (*left*) and ΔZBPs cells (*right*). Epifluorescence imaging of FtsZ-mNeonGreen induced with 20 μM IPTG for 2 hours. Strains: control: bAB219, ΔZBPs: bGS308. F Distances between neighboring Z rings in ΔZBPs cells. Dashed line: estimated spacing between Z rings in non-dividing *B. subtilis* cells, based on cell length (see supplemental methods). Strain bGS308. G Average intensity projections of Z rings in control and ΔZBPs cells (*top*), created by averaging N>800 Z ring images for each strain, and their intensity profiles (*bottom);* shaded areas indicate standard error. Strains: control: bAB219, ΔZBPs: bGS308. Scale bars: 2 μM.

Next, we examined how the ZBPs in combination affect FtsZ dynamics. While none of the ZBPs are individually essential, Δ*sepF* and Δ*zapA* are each synthetically lethal with Δ*ezrA* (*2*). We created a ΔZBPs strain that lacked all ZBPs by knocking out *sepF* and *zapA* and depleting *ezrA* using a xylose-inducible promoter. We depleted EzrA for 7 hours, at which point cells were filamented, indicating that division was blocked. We additionally repeated this for all other synthetically lethal combinations of ZBPs (Fig S5). In all cases, FtsZ treadmilling velocity and subunit lifetime were unchanged from the control (Fig 3D, S6, Movie S6). We note that, although Δ*ezrA* cells have longer FtsZ subunit lifetimes, the lifetimes under these synthetic lethal conditions are statistically indistinguishable from the control. This suggests that the process that elongates FtsZ filaments in the absence of EzrA does not occur under these synthetic lethal conditions, i.e., in the absence of division and when cells are filamented (Supplementary Text 1). Regardless, together these data indicate that ZBPs do not affect FtsZ treadmilling *in vivo*.

Next, we investigated whether the ZBPs instead mediated filament bundling. ZBPs have been shown to mediate FtsZ filament bundling *in vitro* (*19, 21, 23–27*), and lateral interactions between FtsZ filaments have been proposed to play a functional role in cytokinesis (*28, 29*). We, therefore, investigated how each ZBP knockout, individually and in combination, affected Z ring morphology. Overall Z ring morphology was normal in single ZBP knockouts, in the only viable double knockout (Δ*sepF*Δ*zapA*), and in all overexpression conditions except EzrA (Fig S2-4, S7).

However, in the absence of synthetically lethal combinations of ZBPs, cells showed severely altered Z rings: filaments no longer condensed, instead forming loose, regularly-spaced bands occupying regions ~1.6x as wide as control Z rings (Fig 3E-G, S5-6). These bands resemble the transient FtsZ structures that occur prior to Z ring condensation (Fig 4A). Normally, these structures condense into Z rings prior to cell division (Fig 4B), but without ZBPs, condensation never occurs (Fig 3E,G). These results agree with previous observations that ZapA and SepF promote FtsZ bundling, whereas EzrA has previously been described as an inhibitor of Z ring formation and bundling (Supplementary Text 1). Here, we find that the ZBPs work collectively to promote Z ring condensation. Thus, without ZBPs, FtsZ filaments treadmill normally and localize correctly, but cannot condense into Z rings or divide the cell.

**Fig. 4:**
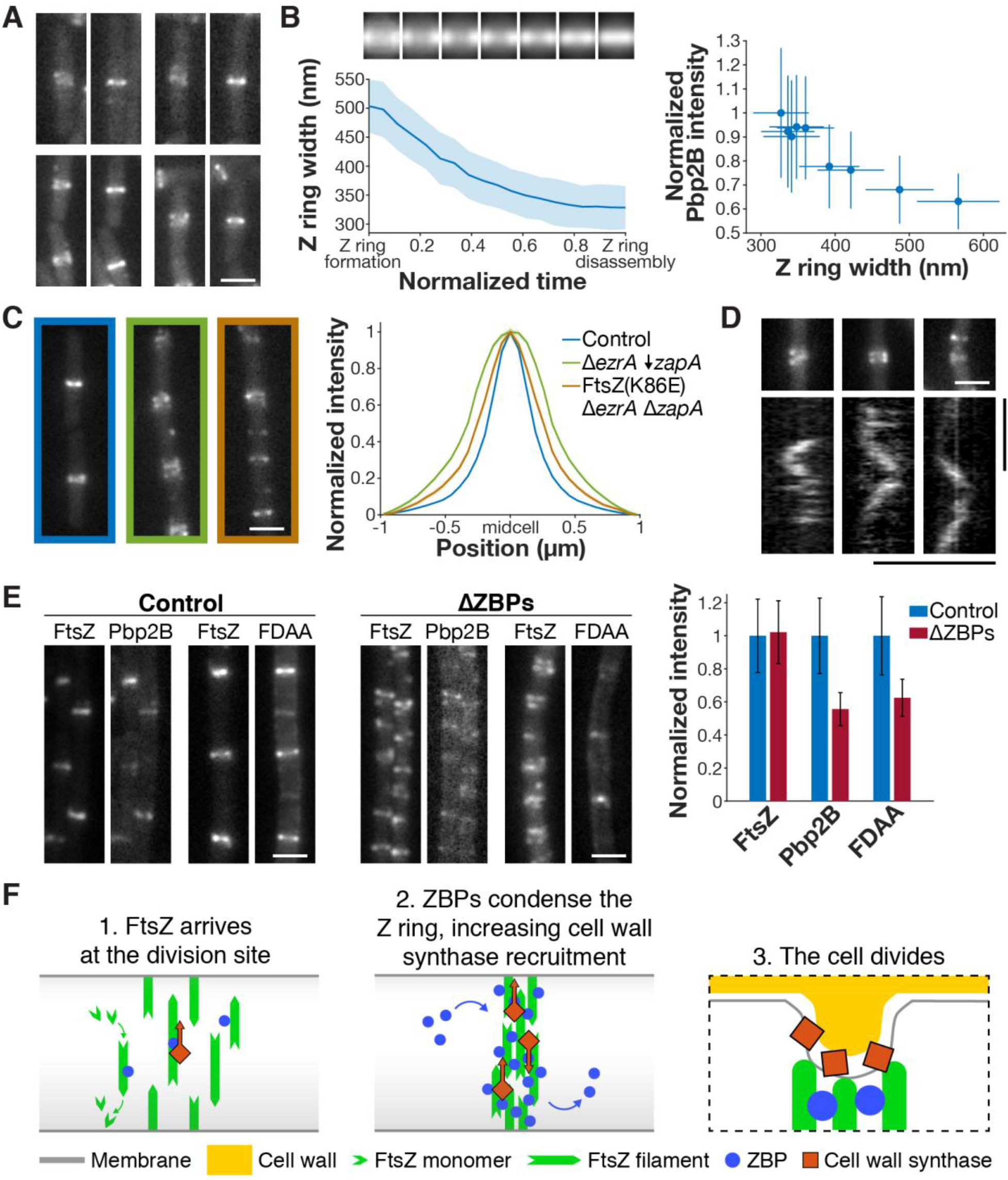
Z ring condensation is required for cell division. **A** The early Z rings before and after condensation in control cells. Each pair of images shows a newly formed Z ring that has not yet condensed (left), and the same Z ring after condensation (right). Z rings were visualized by epifluorescence imaging of FtsZ-mNeonGreen induced with 20 μM IPTG for 2 hours. Strain bAB219. **B** *Left*: Z ring condensation during the cell cycle. *Top*: Average intensity projections of Z rings over the cell cycle, created by averaging Z ring images from each normalized time point. *Bottom*: Z ring width over the cell cycle, measured as the full width at half maximum of the average intensity projections. Time from Z ring formation to Z ring disassembly (defined as the first and last frames in which the Z ring could be detected by peak finding) was normalized for each cell. Shading: bootstrapped standard error. Strain bAB219. *Right*: Pbp2B intensity increases as Z rings condense. Z rings were labeled with FtsZ-HaloTag, induced with 20 μM IPTG for 2 hours and labeled with 5 nM JF549-HTL; for Pbp2B, a Pbp2B-mNeonGreen fusion was used. Error bars: bootstrapped standard error. Strain bGS104. **C** The FtsZ(K86E) mutant rescued the Δ*ezrA* Δ*zapA* synthetic lethal condition and partially restored Z ring morphology, as seen in representative Z ring images (*left*) and intensity profiles (*right*). ↓ indicates depletion. Strains: control: bAB219, Δ*ezrA* ↓*zapA*: bGS293, FtsZ(K86E) Δ*ezrA* Δ*zapA*: bGS463. **D** Pbp2B dynamics in ΔZBPs. Kymographs were drawn at the Z rings indicated above. TIRFM of Pbp2B-HaloTag labeled with JF549-HTL. Strain bMH443 **E** *Left*: Colocalization of Pbp2B with FtsZ and colocalization of FDAA labeling with FtsZ in control cells, visualized by epifluorescence imaging. FtsZ: FtsZ-HaloTag was induced with 20 μM IPTG for 2 hours and labeled with 5 nM JF549-HTL; Pbp2B: Pbp2B-mNeonGreen; FDAA: cells were labeled with 1 mM FDAA for 30 seconds. *Center*: Colocalization of Pbp2B with FtsZ and colocalization of FDAA labeling with FtsZ in ΔZBPs cells. *Right*: Amount of FtsZ, Pbp2B, and FDAA labeling at the division site, measured by fluorescence intensity. Error bars: bootstrapped standard error. Strains for FtsZ and Pbp2B colocalization: control: bGS104, ΔZBPs: bMH445. Strains for FtsZ and FDAA colocalization: control: bMH510, ΔZBPs: bMH508. **F** *Left*: At the start of the cell division process, FtsZ filaments treadmill around the cell circumference at midcell. *Middle*: Stationary ZBPs transiently bind to FtsZ filaments to condense the Z ring, increasing the recruitment of directionally-moving cell wall synthesis enzymes. *Right*: ZBP-driven bundling of FtsZ filaments may also function during cytokinesis, where crowding may induce inward membrane deformations, both concentrating cell wall synthesis to the Z ring and orienting it to divide the cell in two. Scale bars: horizontal: 2 μm, vertical: 1 min.

We next sought to clarify whether the Z ring condensation is specifically due to lateral bundling of FtsZ filaments by ZBPs. If this were the case, we might expect to isolate mutations that promote lateral bundling of FtsZ filaments in cells lacking ZBPs. Thus, we conducted a suppressor screen in the ΔZBPs strain (see Supplemental Methods for details). Whole-genome sequencing of the resulting suppressor candidates revealed a charge-inverting mutation (K86E) in helix H3 of FtsZ; both this helix and the homologous residue have been shown to affect lateral FtsZ filament interactions in *E. coli* (*29, 30*). We hypothesized that this mutation might restore viability in the absence of ZBPs by enhancing filament interactions. Indeed, FtsZ(K86E) restored viability and partially restored Z ring condensation in Δe*zrA* Δ*zapA* cells (Fig 4C, S8).

Interestingly, the FtsZ(K86E) suppressor mutant can rescue the Δe*zrA* Δ*zapA* cells but not other synthetic lethal combinations. Although the ZBPs work collectively to bundle FtsZ filaments, they may each affect bundling differently. Beyond their role as bundlers, the ZBPs have been shown to have distinct functions (*2*). Thus, the fact that FtsZ(K86E) can replace EzrA and ZapA but not SepF may reflect that each ZBP has different effects on FtsZ superstructure.

Finally, we investigated how the bundling of FtsZ filaments by ZBPs affected septal cell wall synthesis, which is required for cell division. Thus, we removed ZBPs and investigated the localization and motion of the division-specific cell wall synthesis enzyme Pbp2B, as well as septal cell wall synthesis activity (4D-E). Pbp2B recruitment to the Z ring decreased by 50% in ΔZBPs relative to control cells (Fig 4E); this is consistent with decreased Pbp2B recruitment at decondensed Z rings in control cells (Fig 4B). We found that in ΔZBPs, Pbp2B nevertheless moved directionally at midcell; because the directional motion of Pbp2B reflects its activity, this suggests that it remains active under these conditions (Fig 4D, S9). To more directly assay the activity of cell wall synthesis enzymes, we measured the incorporation of fluorescent D-amino acids (FDAAs) into the division site (*5*); FDAA incorporation was still present in ΔZBPs but reduced 40% relative to the control (Fig 4E, S9). Thus, septal cell wall synthesis enzymes are still active in the absence of the ZBPs, but Z ring condensation enhances their recruitment to FtsZ filaments at the division site.

Combined with past work, these experiments provide new insights into the mechanisms underlying bacterial cell division. The cell division process begins with short treadmilling FtsZ filaments that are restricted to midcell by negative regulators. Our data reveal that FtsZ filaments treadmill at their biochemical steady state; their dynamics are not modulated by other factors. However, FtsZ cannot form a functional Z ring on its own: ZBPs are also required to bundle FtsZ filaments into a condensed Z ring, transiently interacting with stationary FtsZ subunits without affecting filament dynamics. Z ring condensation increases the recruitment of cell wall synthesis enzymes to the division site, which move around the division site as part of a directionally-moving complex. This condensation is ultimately necessary for cell division (Fig 4F).

These results also yield new insights into the role of Z ring condensation in bacterial cytokinesis. Why is FtsZ filament bundling required for division, and what role does it play in the process? In contrast to previous models (*29, 31*), FtsZ bundling does not modulate FtsZ treadmilling dynamics, but rather condenses the Z ring. While condensation is not required for the activity of division-associated cell wall synthesis enzymes, it may be necessary to concentrate their activity in a small enough region to allow for productive septation. It is also possible that lateral filament association serves to inwardly deform the membrane. FtsZ has been seen to deform liposomes when filaments coalesce (*32, 33*), and crowding of membrane-associated proteins is sufficient to deform membranes (*34*). Such deformations may be easier if the periplasm is iso-osmotic with the cytoplasm (*35, 36*), and therefore the force required for membrane deformation is small. This membrane deformation could promote late protein localization, or the late proteins might bind more readily to the early proteins once they have condensed; either mechanism could explain the enhancement of Pbp2B recruitment by Z ring condensation. This membrane deformation could then direct circumferential septal wall synthesis inward to divide the cell (*5, 37*).

## Acknowledgements

We thank the Garner lab, especially A. Bisson-Filho, Y. Sun, and S. Wilson, for discussions, and L. Lavis for JF dyes. TIRF-SIM was performed at the Advanced Imaging Center at the Janelia Research Campus, a facility jointly supported by the Gordon and Betty Moore Foundation and Howard Hughes Medical Institute. Funding: This work was funded by National Institutes of Health Grants DP2AI117923-01 to ECG, as well as support from the Volkswagen Foundation, NSF GRFP (DGE1144152) to GRS, and the Physiology course at the Marine Biological Laboratory at Woods Hole. This work was supported by the NSF-Simons Center for Mathematical and Statistical Analysis of Biology at Harvard (1764269) and the Harvard Quantitative Biology Initiative. Author Contributions: GRS, MJH, and ECG designed experiments. GRS and MJH conducted experiments and analyzed data, with help from SB, BP, JR, VY on initial optimization. GRS, MJH, and ECG wrote the manuscript. Competing Interests: Authors declare no competing interests. Data and materials availability: Code is available at bitbucket.org/garnerlab/squyres-2020.

