## Supplementary Materials for "Dynamics of bacterial cell division: Z ring condensation is essential for cytokinesis"

### Materials and Methods

#### Culture growth

Strains were stored as glycerol stocks at -80˚C. At the start of each experiment, strains were streaked onto LB agar plates containing the appropriate antibiotic and incubated overnight at 37˚C. For strains whose survival was dependent on the induction of a promoter, these plates were additionally top spread with xylose or IPTG at the appropriate concentration. For imaging, single colonies were inoculated into 1 mL casein hydrolysate (CH) media and grown on a roller at 37˚C until they reached mid-exponential-phase growth ($OD_{600}$ around 0.2). Cells were back diluted 1:10 and again grown until mid-exponential phase; this process was repeated until cells were ready for imaging. Alternately, cells were grown overnight in CH on a roller at 25˚C. These cultures were grown in a 1:10 dilution series out to 1:10,000; the next day, the culture whose $OD_{600}$ was nearest to 0.2 was back diluted 1:10 and grown in CH at 37˚C as above.

#### Microscopy: sample preparation

Cells were grown in 1 mL CH media at 37˚C to $OD_{600}$ around 0.2 as described above. Cultures were concentrated approximately 10-fold by centrifugation for 2 minutes at 7,000 RPM and resuspended in CH. Agarose pads were prepared using square plastic frames with inner dimensions 1.5 cm x 1.5 cm x 1 mm. Frames were placed on a cleaned glass pane, molten CH + 2% agarose was poured into the frames, and a second glass plane was placed on top to form a mold. Pads were allowed to solidify at room temperature, and excess agarose was cut away from the outside of the frame. To prepare slides for imaging, 2 µL of concentrated cells were pipetted onto a base-washed coverslip, and an agarose pad was placed on top. For multi-hour acquisitions, glass-bottom dishes (MatTek) were used instead of coverslips; these were also base-washed before use, and a moist KimWipe was wound around the edge of the dish to retain humidity.

#### Microscopy: phase contrast, epifluorescence, and TIRFM

Phase contrast, epifluorescence, and Total Internal Reflection Fluorescence Microscopy (TIRFM) images were collected on a Nikon Ti-E microscope using a Nikon CFI Plan Apochromat DM Lambda 100X Oil objective, 1.45 NA, phase ring Ph3. Cameras used were an ORCA-Flash4.0 V2 sCMOS (Hamamatsu) and an iXon Ultra 897 EMCCD (Andor). Fluorescence excitation was achieved using a MLC4008 laser launch (Agilent) with 405 nm, 488 nm, 561 nm, and 647 nm lasers. For fluorescence emission, a C-NSTORM QUAD filter cube was used, along with an additional ET525/50m filter for green emission and ET600/50m filter for red emission (Chroma). The microscope was enclosed in a chamber heated to 37˚C.

#### Microscopy: SIM-TIRF

Live-cell SIM data were acquired as described previously (*38*) on a Zeiss Axio Observer.Z1 inverted microscope outfitted for structured illumination. An Olympus ×100/1.49NA objective was used instead of the Zeiss 1.45 NA objective because the slightly larger NA of the Olympus objective gives higher tolerance for placing the excitation beams inside the TIRF annulus. Data were acquired at 1 s frame rates for 2 minutes with 20 ms exposures from a 488 nm laser for each rotation. TIRF-SIM images were reconstructed as described previously (*38*).

#### Induction, depletion, and HaloTag labeling

For FtsZ imaging, FtsZ-mNeonGreen or FtsZ-HaloTag were expressed from the IPTG-inducible pHyperSpank promoter. In all cases, FtsA was co-expressed from the same promoter, preserving the native operon structure. These constructs were merodiploid, meaning that the inducible FtsZ constructs were cloned into the chromosome at an ectopic site; the native untagged operon remained intact. Labeled FtsZ was induced by adding 20 µM IPTG to the growth media for 1 hour before imaging. For single-molecule imaging, strains containing FtsZ-HaloTag were labeled by adding 20 pM JF549-HaloTag Ligand (JF549-HTL) to the growth media for 1 hour before imaging (*7*). For total labeling, 5 nM JF549-HTL was used. For overexpression, the FtsZ-mNeonGreen construct was induced with 100 µM IPTG for 1 hour before imaging.

Single-molecule imaging of other divisome proteins was conducted as follows. FtsA, EzrA, and ZapA HaloTag constructs were expressed as a sole copy from their native promoters and labeled with 50 pM, 300 pM, and 600 pM JF549-HTL, respectively. SepF-HaloTag was expressed as a merodiploid under an IPTG-inducible promoter; no IPTG was added, as leaky expression from the promoter was sufficient for single-molecule imaging. SepF-HaloTag was labeled with 200 pM JF549-HTL. DivIB, DivIC, and FtsW HaloTag constructs were expressed as sole copies from xylose-inducible promoters. They were induced with 1 mM, 5 mM, and 8 mM xylose, and labeled with 400 pM, 500 pM, and 300 pM JF549-HTL, respectively. FtsL-HaloTag and Pbp2B-HaloTag were expressed as a sole copy from an IPTG-inducible promoter, induced with 30 µM and 20 µM IPTG, and labeled with 400 pM and 200 pM JF549-HTL, respectively. All JF549-HTL labeling was performed for 15 minutes before imaging; when JF549-HTL concentrations were sufficiently high, cells were washed once in 1 mL CH media before imaging to remove excess dye.

For overexpression of ZBPs, xylose was added at the indicated concentration for 2 hours before imaging. For depletion of ZBPs, cells were grown initially in 1 mM xylose; xylose was then withdrawn, and cells were imaged at the point when they had filamented but were still alive, approximately 7 hours after xylose withdrawal. For imaging of Pbp2B dynamics in these mutants, cells were grown overnight in 100 µM IPTG and 1 mM xylose; 7 hours before imaging, xylose was withdrawn and the concentration of IPTG was reduced to 20 µM. Pbp2B-HaloTag was labeled with 100 pM JF549-HaloTag Ligand for 15 minutes before imaging.

**Velocity measurements**

To measure FtsZ treadmilling velocity, cells expressing FtsZ-mNeonGreen were imaged by TIRFM. Time lapses were taken using the sCMOS camera with 1 s exposures for 4 minutes total; after each time lapse, a phase-contrast image was taken to visualize cells. Velocity was measured by kymograph analysis as in (*5*). Kymographs were created from these time lapses of fluorescently labeled FtsZ filaments by manually drawing ROIs along the short axis of cells in ImageJ. Regions of these kymographs containing diagonal bands of fluorescence, representing directional treadmilling, were selected and their slopes were measured manually in ImageJ.

Velocity measurements of the single-molecule motions of DivIB, DivIC, FtsL, FtsW, and Pbp2B were taken similarly; cells were labeled for single-molecule imaging as described above and imaged by TIRFM. Each of these cells additionally expressed FtsZ-mNeonGreen to visualize the division site. Time lapses were taken using the sCMOS camera with 1-second exposures for 2-4 minutes total; before and after each time lapse, a phase-contrast image was taken to visualize cells, and a green epifluorescence image was taken to visualize the division site. Kymograph analysis of velocities was performed as summarized above; in this case, molecules that colocalized with the division site were specifically selected for analysis. A summary of these results is provided in Table S3.

To characterize the stationary behavior of EzrA, SepF, and ZapA, cells were labeled for single-molecule imaging and imaged by TIRFM as above. These cells also expressed FtsZ-mNeonGreen to visualize the division site. The microscopy protocol was identical to that in the previous paragraph; molecules that colocalized with the division site were selected for analysis.

**Cell segmentation**

Phase-contrast images of cells were segmented using DeepCell, a deep learning-based cell segmentation platform (*39*). A different custom net was trained for each combination of objective and camera used. Training sets were manually generated and varied in size but were generally around 20 images each. For cells in synthetic lethal conditions, masks were refined manually to omit dead cells.

#### Custom MATLAB software

Custom code is available at https://bitbucket.org/garnerlab/squyres-2020

#### Single-molecule lifetime measurements

To measure the single-molecule lifetimes of FtsZ and the ZBPs, HaloTag fusions were grown and labeled for single-molecule imaging as described above. TIRFM time lapses were taken using the emCCD camera, with 500 ms exposures for 4 minutes total; after each time lapse, a phase-contrast image was taken to visualize cells. To analyze the data, first, the phase images of cells were segmented using DeepCell to generate cell masks. Next, TrackMate was used to identify single particles in the image and preliminarily link them together (*40*). Spots were detected with a 1.5-pixel radius and an intensity threshold that was manually selected for each data set. Spots were then linked roughly into tracks, with a 3-pixel linking distance and a maximum gap of 10 frames; in this way, only localizations with at least one other spot detected nearby were considered for further analysis, which decreased computational load in the next stage. These data were exported into MATLAB for further analysis.

The track list from TrackMate was then filtered and converted to intensity traces. First, the spot positions in each track were averaged to generate a mean position of each spot. Next, spots that were not inside cells were excluded using the cell masks generated by DeepCell. Spots within 3 pixels of one another were then combined, and a new average position was calculated, weighted based on the length of each track. Then, for each spot, an intensity trace was generated: intensity was averaged in a 5 x 5 pixel window around the mean spot position, and the local background was averaged in a 2-pixel frame around the window and subtracted to generate a background-subtracted trace. Finally, intensity traces were filtered; only traces with a maximum background-subtracted intensity above 500 counts were included for further analysis.

To measure each single molecule lifetime, these intensity traces were fit to a Hidden Markov Model (HMM) using the MATLAB package vbFRET (*41*). To rule out spots which contained no single-molecule fluorescence events, and to exclude cases where multiple single molecules overlap, models were fit with 1, 2, 3, and 4 states. Bayesian model selection was used to select the best-fitting model, and only traces in which the 2-state model fit best were included. Traces for which the difference between state 1 (no fluorescence) and state 2 (single-molecule fluorescence) was less than 60 counts were also discarded. The duration of each state 2 event was measured; dwell times less than 2 seconds (4 frames) were discarded, as were events that overlapped with the start or end of the trace since they cannot be measured accurately. Traces containing more than 2 events were also excluded. The resulting single-molecule lifetimes were fit to a single exponential distribution and the mean lifetime was computed. To compare lifetimes, p-values were calculated using a Wilcoxon rank-sum test. Results are summarized in Table S4.

#### Single-molecule lifetime measurements: inside versus outside of the Z ring

To compare the single-molecule lifetime of FtsZ inside and outside of the Z ring, cells expressing FtsZ-HaloTag were prepared for imaging as described previously. These strains also expressed mNeonGreen-Pbp2B to visualize the division site. TIRFM time lapses were taken using the emCCD camera, with 500 ms exposures for 4 minutes total; before and after each time lapse, a phase-contrast image was taken to visualize cells, and a green epifluorescence image was taken to visualize the division site. To compare single-molecule lifetimes inside and outside of the Z ring, ROIs were manually drawn around each division site, as seen in Figure 2D; lifetimes were analyzed as described above and assigned to be inside or outside of the Z ring based on the drawn ROIs. 38% of single molecules were identified as being inside Z rings, consistent with quantitative fluorescence measurements in *B. subtilis* that found 30-35% of FtsZ in the Z ring (*13*).

**Single-molecule lifetime measurements: manual**

To confirm that the automated single-molecule lifetime measurements described above were accurate, single-molecule lifetimes were also measured by hand. FtsZ-HaloTag was imaged as described in the Single-molecule lifetime measurements section above, although the sCMOS camera was used instead of the emCCD camera for ease of visualization. Kymographs were drawn manually in ImageJ. These kymographs were then examined for single-molecule events, and the duration of these events was measured manually in ImageJ.

**Z ring identification, spacing, width, and average projections**

To visualize the Z ring, cells expressing FtsZ-mNeonGreen under an IPTG-inducible promoter were grown for imaging as usual. Cells were imaged using the sCMOS camera; at each position, one phase-contrast image to visualize cells and 1 green epifluorescence image to visualize Z rings were taken. To identify Z rings in the image, cells were segmented using DeepCell to generate binary masks. The pill mesh function in Morphometrics was then used to generate midlines down the long axis of each cell (*42*). Using custom MATLAB code, the fluorescence intensity of FtsZ was averaged along each cell midline by taking the average along each mesh spline, these intensity traces were smoothed, and Z rings were identified by peak detection.

To measure the spacing between Z rings, the distances between neighboring peaks were measured. The predicted spacing between Z rings was calculated as follows. Because ΔZBPs cells do not divide, Z rings will not disassemble once they are formed, and Z rings assemble around 25% of the way through the cell cycle (*3*). Thus, the expected Z ring spacing is equal to the cell length for cells at or below the 25th percentile for length and is ½ the cell length for cells above the 25th percentile. Cell length data for WT cells in CH media were obtained from (*43*).

To plot intensity traces of Z rings, a 1 µm region around each peak was sub-selected from each intensity trace, and these traces were averaged to create an average intensity trace. The Z ring width was measured by calculating the full width at half maximum of the Z ring peak in these intensity traces. Results are summarized in Table S5. To display average projections of Z rings, regions of each corresponding cell were sub-selected around each peak, using the meshes to align and straighten cells and normalize cell width; these images were averaged to create an average projection.

**Z ring width across the cell cycle**

To quantify the appearance of the Z ring over the cell cycle, FtsZ-mNeonGreen cells were grown as above. Cells were then imaged in phase and epifluorescence as above, repeated every minute for 2 hours. Time lapses were registered in ImageJ, and phase images were segmented using DeepCell. Morphometrics was used to generate midlines down each cell, as well as to track cells over time. Fluorescent images and cell meshes were then imported into MATLAB for further analysis.

First, cell tracks from Morphometrics were filtered for quality control. Cell tracks with a duration less than 20 frames (20 minutes) were discarded. Additionally, the total cell length for each frame in the track was then fit to a line, and tracks for which the $R^2$ of this fit was less than 0.99 were also discarded. Next, Z rings in the cell in each frame were identified as described above. These Z rings were then linked together between frames by particle tracking to create Z ring tracks: Z rings were linked if they were within 5 pixels (325 nm) and 5 frames (5 minutes) of one another. Only Z ring tracks between 20 and 40 minutes in duration were considered for further analysis.

Time was normalized for each track, and Z ring width and average projections were computed as described above.

**FtsZ and Pbp2B colocalization**

To quantify the colocalization between FtsZ and Pbp2B at the division site, cells were grown containing FtsZ-HaloTag under an IPTG-inducible promoter and Pbp2B-mNeonGreen under its native promoter. The ZBP depletion strain was depleted for 7 hours before imaging as usual. FtsZ-HaloTag was induced with 20 µM IPTG and labeled with 5 nM JF549 for 1 hour before imaging. Cells were imaged at 20 positions using the sCMOS camera; at each position, in order, 1 phase contrast image, 1 red epifluorescence image, and 1 green epifluorescence image were taken. Images were background-subtracted in ImageJ with rolling ball radius 50. To analyze, Z rings were identified in each cell as described above. These same peak regions were selected from the corresponding Pbp2B image, to visualize Pbp2B intensity at the Z ring. Finally, the area under these peak regions was calculated to estimate the amount of FtsZ and Pbp2B at midcell in each strain.

**Cell wall synthesis labeling**

For live-cell fluorescent D-amino acid (FDAA) labeling of cell wall synthesis, cells were grown for imaging as normal; the ZBP depletion strain was depleted for 7 hours before imaging as usual. To visualize the division site, these cells also expressed FtsZ-HaloTag, which was induced with 20 µM IPTG and labeled with 5 nM JF549 for 1 hour before imaging. Cells were pelleted at 8000 RPM for 30 seconds and resuspended in 10 µL CH + 1 mM fluorescent D-lysine (FDL). Cells were incubated for 3 minutes to allow labeling to occur, after which 1 mL CH was added to the tube to halt FDL labeling. Cells were pelleted at 8000 RPM for 30 seconds, resuspended in 100 µL CH, and immediately placed under an agarose pad for imaging. The average time between the end of FDL labeling and the start of imaging was 3 minutes and 20 seconds. Image acquisition was automated to increase speed and took an additional 1 minute. Cells were imaged at 10 positions using the sCMOS camera; at each position, in order, 1 phase contrast image, 1 red epifluorescence image, and 1 green epifluorescence image were taken.

Images were background-subtracted in ImageJ with a rolling ball radius of 50. To analyze colocalization, Z rings were identified in each cell as described above. These same regions were selected from the corresponding FDAA image. To correct for differences in labeling efficiency between cells, the FDAA signal at midcell was normalized to signal in the nearby sidewall. Finally, the area under these peak regions was calculated to estimate the amount of FtsZ and FDAA at midcell in each strain.

**Pbp2B dynamics**

To visualize Pbp2B dynamics at the division site, cells expressed both Pbp2B-HaloTag from an inducible promoter to visualize single-molecule Pbp2B dynamics and FtsZ-mNeonGreen from an ectopic site under its native promoter to visualize the Z ring. Cells were plated overnight on LB plates top spread with 100 µM IPTG; 1 mM xylose was also added to the plate for the synthetic lethal depletion strain. The following day, single colonies were inoculated into 1 mL CH + 1 mM xylose + 100 µM IPTG cultures and grown overnight at room temperature with shaking. The next morning, cells were washed once in 1 mL CH media and then grown for 7 hours in CH media + 20 µM IPTG without xylose. This both began the depletion process for the synthetic lethal strain and decreased Pbp2B expression to a level suitable for single-molecule analysis. 15 minutes before imaging, cells were labeled with 100 pM JF549-HaloTag Ligand. Cells were imaged by TIRFM: time lapses were taken using the sCMOS camera with 1-second exposures for 4 minutes total; before and after each time lapse, a phase-contrast image was taken to visualize cells, and a green epifluorescence image was taken to visualize the division site. Kymographs were created manually as described above. Z ring images were assigned to these kymographs by extracting a 61 x 61 pixel region from the FtsZ epifluorescence image taken before TIRFM, centered on the midpoint of the kymograph.

To characterize Z rings at which Pbp2B moved directionally, directionally moving Pbp2B particles were identified by particle tracking. Particles were tracked using TrackMate with the following parameters: spots with a diameter of 400 nm were identified using the Laplacian of Gaussians (LoG) detector; these spots were tracked over a 100 nm search radius using the Sparse LAP Tracker with no frame gaps. The resulting tracks were further filtered to obtain a selection of tracks with clear directional motion using a custom MATLAB script, as described previously (*5*). Tracks between 10 and 25 seconds and with end-to-end displacement above 225 nm were included for further analysis; further quality control was achieved by selecting tracks well fit ($r_2$ >0.95) linearly to a log-log plot of mean squared displacement (MSD) vs time interval (Δt). Any remaining diffusing particles were omitted by ensuring a nonzero velocity from the fit $MSD(\Delta t) = 4*D*\Delta t + (v*\Delta t)_2$ (v: velocity; D: diffusion constant). Tracks were assigned to cells using phase images segmented as described above. Z rings were identified in these cells (described above) and tracks were assigned to the nearest Z ring up to a maximum distance of 1 µm.

**Suppressor screen**

To isolate potential mutations that could suppress the synthetic lethality of the ZBPs, cells of strain bGS308 (Δ*sepF*, Δ*zapA*, xylose-inducible *ezrA*) were plated overnight on LB plates without xylose. Colonies grew overnight, as expected since EzrA depletion is slow and cells that are inhibited for division can grow for some time before death. The following day, colonies were inoculated into 3 mL LB cultures in triplicate and grown for 8 hours at 37˚C with shaking. During the 8 hours of growth, the OD first increased, then decreased as cell death occurred, and then began to increase again. After 8 hours, 200 µL of each culture was plated on LB plates and grown overnight. The following day, 5 colonies from each plate were restruck for single colonies. Each colony was also patched onto LB plates top-spread with 25 mM xylose: because EzrA overexpression is lethal, candidates with an intact xylose-inducible promoter were expected to die on high xylose plates. Of the 15 candidate colonies, 4 did not grow after restreaking, and 3 had become insensitive to high xylose; the remaining 8 were submitted for whole-genome sequencing.

All 8 candidates had mutations in the xylose-inducible promoter; 4 candidates also had additional mutations. Of these, strain bGS390 (containing the FtsZ(K86E) mutation) was selected for further analysis. To verify that this mutation was capable of suppressing ZBP synthetic lethality, the FtsZ(K86E) mutation was introduced into a WT background. Individual Δ*ezrA*, Δ*sepF*, and Δ*zapA* mutations were introduced into this background and combined by crossing. As expected, the Δ*sepF* Δ*zapA* double mutant was viable, since these proteins are not synthetically lethal even in the WT background. Additionally, the normally synthetically lethal Δ*ezrA* Δ*zapA* mutants could be combined in the FtsZ(K86E) background, verifying that this mutation is a bona fide suppressor; the presence of the FtsZ(K86E) mutation was confirmed in this strain by Sanger sequencing. As a control, a Δ*ezrA* Δ*zapA* cross was attempted in parallel in the WT background; as expected, although colonies appeared on the transformation plate after overnight incubation, these colonies could not be further grown in liquid culture and became transparent after an additional day of incubation, verifying that these mutations are indeed synthetically lethal.

To rule out the possibility of additional suppressors arising during cloning, a FtsZ(K86E) Δ*zapA* xylose-inducible *ezrA* strain was constructed. This strain was maintained in 1 mM xylose during the cloning process, conditions under which cells containing WT FtsZ are viable; this same process was used to generate the synthetic lethal depletion mutants. After cloning, xylose was withdrawn; unlike WT FtsZ-containing cells, which die after xylose is withdrawn, FtsZ(K86E) mutant cells remained viable after xylose was withdrawn.

Interestingly, neither the Δ*ezrA* Δ*sepF* nor the Δ*ezrA* Δ*sepF* Δ*zapA* mutants could be constructed in the FtsZ(K86E) background. We suspect that the ability of this mutant to survive in the ↓*ezrA* Δ*sepF* Δ*zapA* condition during the initial suppressor screen was due to some leaky expression from the xylose promoter, which may have been enhanced by the mutations in the promoter that arose during the screen.

**Strain construction**

All strains were constructed in *B. subtilis* strain PY79; strains used in this study are listed in Table S1. Constructs were assembled by PCR amplification and Gibson cloning. These Gibson products were transformed directly into competent *B. subtilis*, where they were integrated into the chromosome by homologous recombination with homology regions that were engineered at each end of the construct. Transformants were selected by growth on LB plates containing the appropriate antibiotic. The resulting strains were verified by PCR and, when appropriate, by sequencing. Constructs used in this study, as well as the plasmids used to create each construct, are listed in Table S2.

To combine constructs in the same strain, parent strains containing the constructs to be combined were crossed by transforming genomic DNA from one strain into the other. When two strains to be combined contained the same antibiotic marker, the marker was removed from one of the parent strains. All antibiotic resistance cassettes used were engineered with loxP sites flanking the cassette, and so these cassettes could be removed by transforming cells with a plasmid that expresses Cre recombinase (plasmid pDR244, a gift from David Rudner). This plasmid also has a temperature-sensitive origin of replication, so after incubating cells at 30˚C for 24 hours to remove the antibiotic cassette, cells were shifted to 45˚C and incubated overnight to remove the plasmid. The removal of the cassette was verified by lack of growth on antibiotic selective plates.

Many of the strains used to investigate synthetic lethal conditions, namely bGS204, bGS206, bGS290, bGS293, bGS297, bGS298, bGS306, bGS308, bGS315, and bGS331, were additionally verified by whole genome sequencing to confirm that no suppressor mutations had arisen during cloning.

### Supplementary Text 1

Understanding EzrA's function has been complicated by EzrA's apparently contradictory effects on the Z ring. On one hand, EzrA has been repeatedly characterized as a negative regulator of FtsZ polymers (*13*, *18*, *21*, *22*, *26*, *44*, *45*). On the other hand, EzrA is known to be synthetically lethal with SepF and ZapA, which are positive regulators of FtsZ (*19*, *46*). It has also been shown that EzrA's role in inhibiting polar Z ring formation and EzrA's role at midcell are separable: mutants in EzrA can be made that disrupt one of these functions but not the other (*47*, *48*).

The results presented here also indicate that EzrA has two separable functions. We find that EzrA plays a positive role in condensing the Z ring *in vivo*, working together with SepF and ZapA to promote lateral bundling of the Z ring. Z ring condensation is essential for cell division, and this function explains the synthetic lethalities of Δ*ezrA* mutants with Δ*sepF* and Δ*zapA* mutants (*19*, *46*). We additionally find that EzrA decreases the length of FtsZ filaments. This is consistent with the inhibition of Z ring formation by *ezrA* overexpression (*45*) and the fact that Z rings in Δ*ezrA* cells recover more slowly after photobleaching (*13*). We find that this length decrease does not change FtsZ's treadmilling dynamics, and thus the concentration of free FtsZ monomer. This agrees with *in vitro* results that EzrA has no effect on FtsZ's GTPase activity (*26*, *45*). Because EzrA has also been shown to increase the amount of FtsZ needed to form polymer (*18*, *21*, *45*), and because EzrA is highly expressed in the cell (10,000-20,000 molecules per cell, versus 5,000 for FtsZ) (*45*, *49*), this length decrease is likely due to monomer sequestration, as previously proposed (*21*).

Our results also indicate that these two functions are separable. In cells missing both EzrA and one or both of its synthetically lethal partners Δ*sepF* or Δ*zapA*, Z ring condensation is disrupted but we no longer observe a change in filament length. There are several possible mechanisms by which EzrA's effect on length may be absent in these conditions. One possibility is that this effect happens specifically during Z ring constriction, as has been previously suggested for EzrA; because cytokinesis is inhibited in these conditions, such effects would be lost (*19*, *50*). More generally, inhibition of cell division may trigger stress responses that affect FtsZ polymer equilibria. Regardless, the observation that EzrA's effects are separable is consistent with previous studies (*47*, *48*), and our results indicate that EzrA's role in bundling is of primary importance for cell division.

**Figure S1: Controls for FtsZ lifetime measurements**

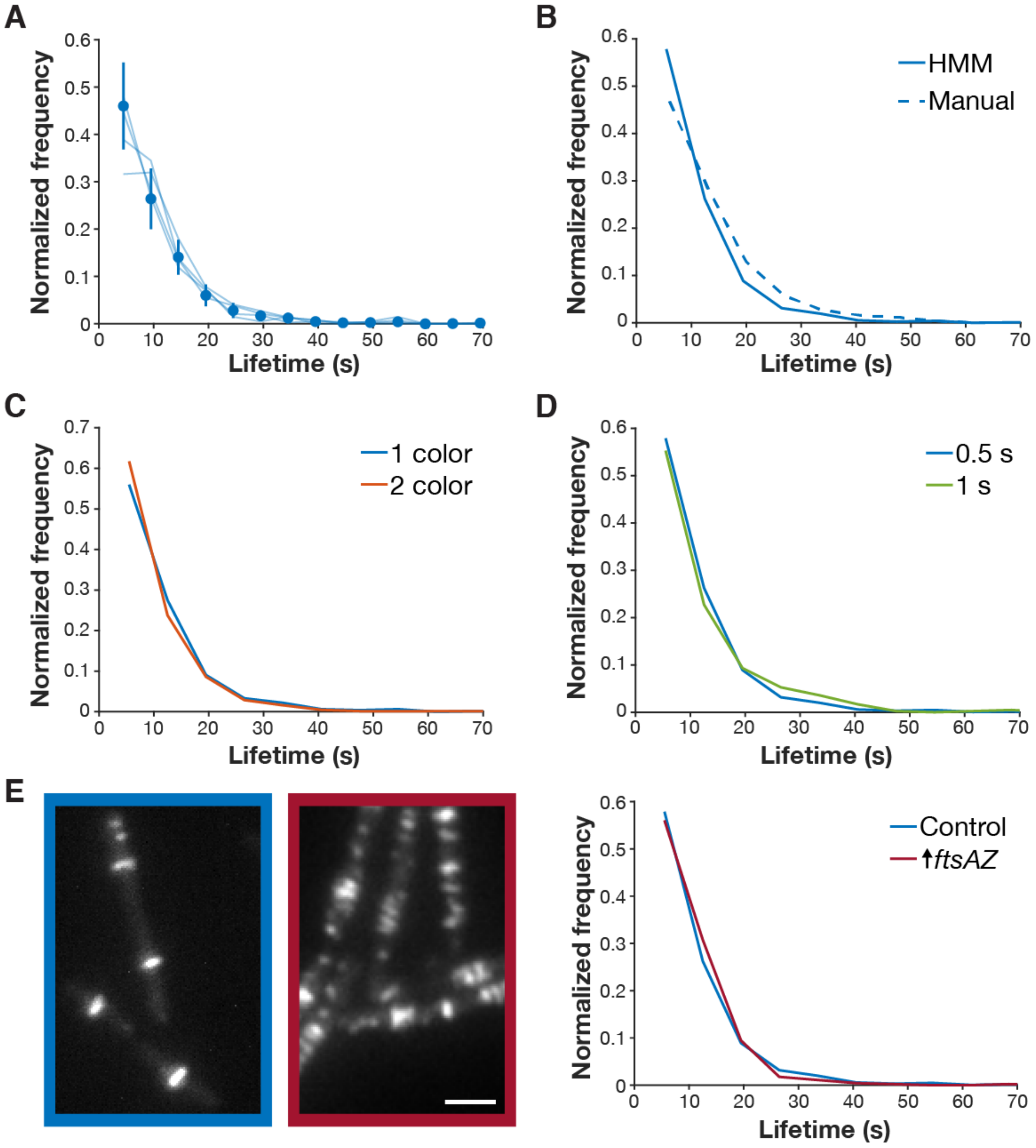


**A** FtsZ subunit lifetime is consistent across experimental replicates. To measure lifetimes, cells expressing FtsZ-HaloTag were induced with 20 µM IPTG for 2 hours, labeled with 40 pM JF549-HTL, and then imaged by TIRFM. Light curves: 4 experimental replicates for which N > 200. Points represent combined data from all experimental replicates (17 replicates). Error bars: weighted standard deviation of distributions for all replicates. Combined data from strains bAB309 and bGS104.

### Figure S1 continued

**B** FtsZ subunit lifetime is consistent across measurement techniques. Lifetime distributions were measured using an automated hidden Markov model (HMM) based analysis pipeline (same data as in A, solid line) and manually for N=265 particles (dashed line).

**C** FtsZ subunit lifetime is not affected by Pbp2B tagging. The 1-color strain (bAB309) contains labeled FtsZ-HaloTag, induced as a second copy with 20 µM IPTG for 2 hours; the 2-color strain (bGS104) contain both this FtsZ-HaloTag construct and a native mNeonGreen-Pbp2B fusion, which was used to localize the division site.

**D** FtsZ subunit lifetime is not affected by photobleaching. If the measurements were affected by photobleaching, the measured lifetimes would increase when we decrease the imaging interval; however, we see that the lifetime distributions are consistent for images taken at 0.5-second intervals and 1-second intervals. For images at 0.5-second intervals, images were acquired continuously with 0.5-second exposures (same data as in A). For images at 1-second intervals, images were acquired with the same settings, with 0.5 seconds of exposure and 0.5-second intervals without illumination (strain bAB309, same method as in A).

**E** Co-overexpression of FtsA and FtsZ increases the number of FtsZ filaments in the cell (*left*) but does not change FtsZ subunit lifetime (*right*). The increased number of filaments that form upon FtsAZ overexpression is consistent with steady-state treadmilling of FtsZ. The fact that the subunit lifetime does not change when FtsAZ is overexpressed further indicates that the additional FtsZ forms new filaments rather than elongating existing filaments. A second copy of *ftsAZ* is expressed from an IPTG-inducible promoter with 100 µM IPTG for 2 hours. Strains, *left*: control: bAB219 induced with 20 µM IPTG, ↑*ftsAZ*: bAB219 induced with 100 µM IPTG. Strains, *right*: control: bAB309 and bGS104 induced with 20 µM IPTG, ↑*ftsAZ*: bAB309 induced with 100 µM IPTG. All inductions in this panel were for 2 hours. Scale bar: 2 µm.

**Figure S2: Effects of individual ZBP knockouts on cell and Z ring morphology**

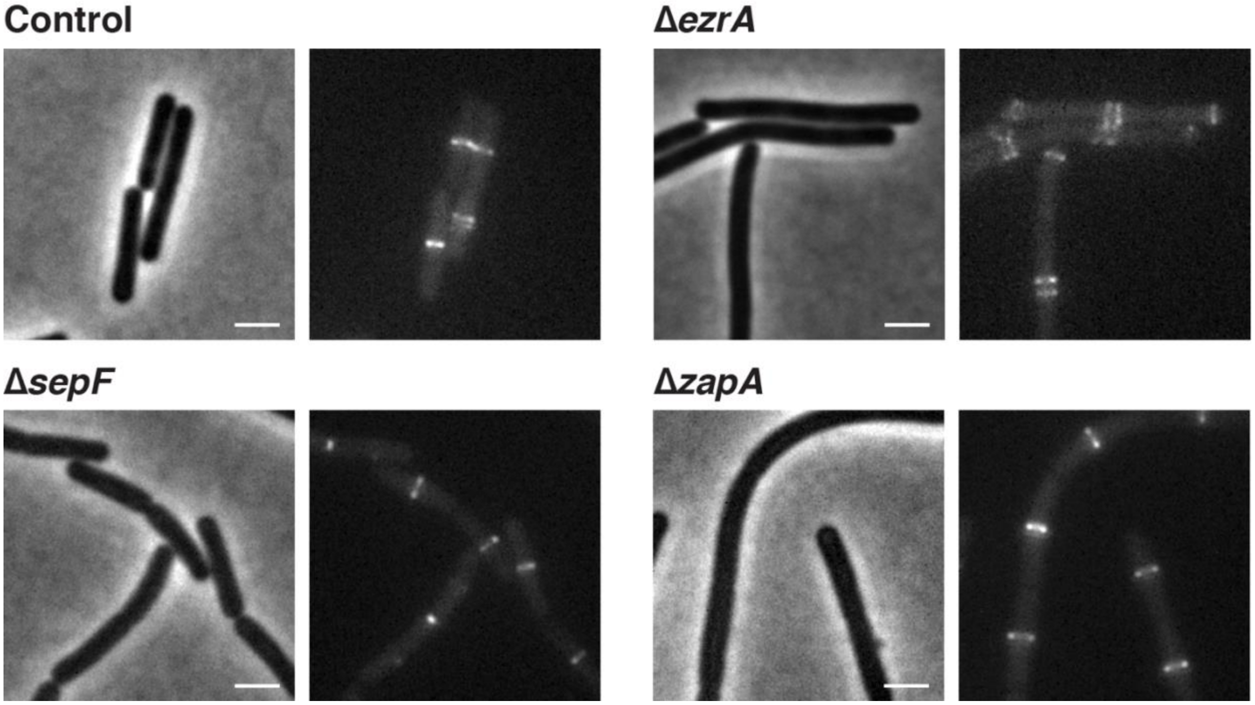


Each pair of images shows cell morphology (phase-contrast imaging, *left*), and Z ring morphology (epifluorescence images of cells expressing FtsZ-mNeonGreen induced with 20 µM IPTG for 2 hours, *right*) in control cells, compared to cells with individual ZBPs deletions. As expected, Δ*ezrA* cells have Z rings near their poles (*18*); Δ*sepF* and Δ*zapA* cells have normal Z rings. Strains: control: bAB219, Δ*ezrA*: bGS256, Δ*sepF*: bGS254, Δ*zapA*: bGS250. Scale bars: 2 µm.

**Figure S3: Effects of SepF and ZapA overexpression**

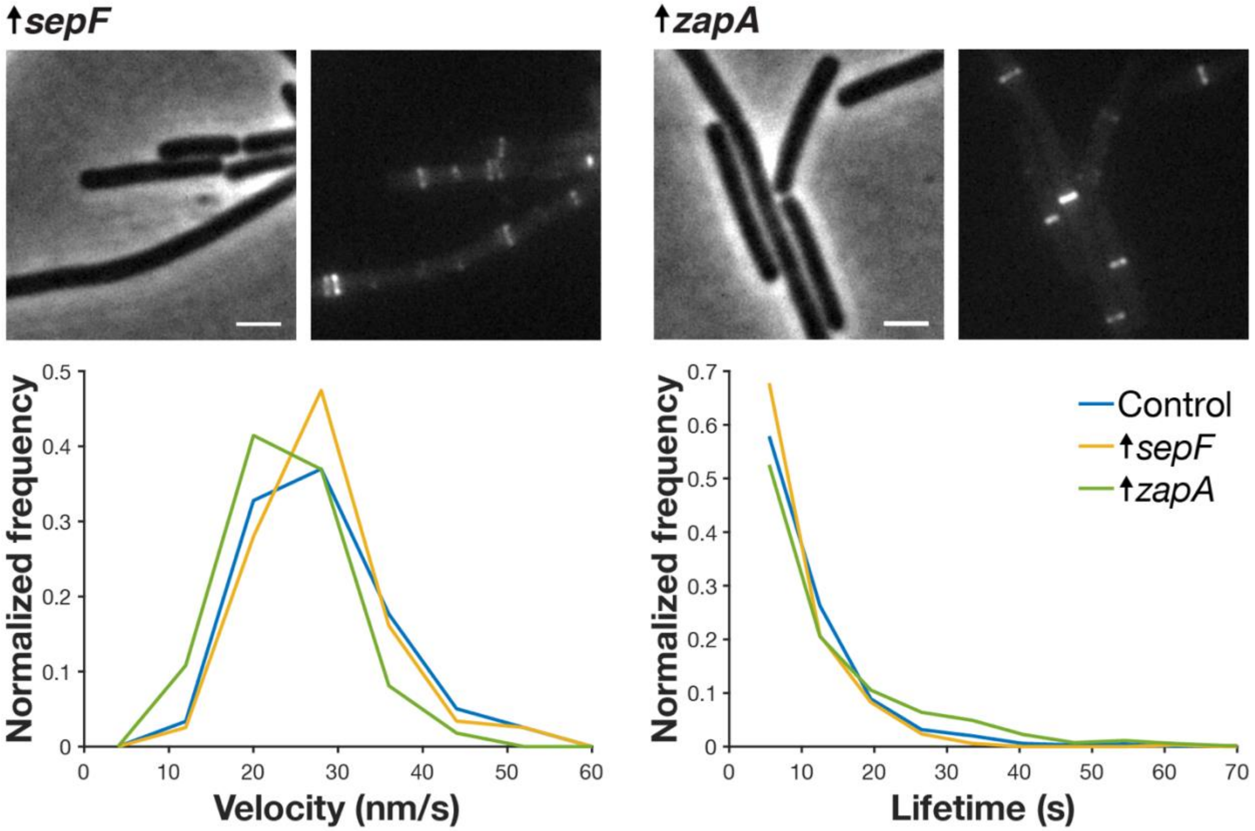


**Top** Each pair of images shows cell morphology (phase-contrast imaging, *left*), and Z ring morphology (epifluorescence images of cells expressing FtsZ-mNeonGreen induced with 20 µM IPTG for 2 hours, *right*), in cells overexpressing SepF and ZapA. These cells have normal Z ring morphology except for some polar Z rings in SepF-overexpressing cells. Second copies of *sepF* and *zapA* were expressed from a xylose-inducible promoter with 30 mM xylose for 2 hours. Strains: ↑*sepF*: bGS260, ↑*zapA*: bGS259. Scale bars: 2 µm.

**Bottom** *sepF*- and *zapA*-overexpressing cells have similar FtsZ treadmilling velocities (*left*) and subunit lifetimes (*right*) to control cells. For velocity measurements, FtsZ-mNeonGreen was induced with 20 µM IPTG for 2 hours, imaged by TIRFM, and analyzed from kymographs. For lifetime measurements, FtsZ-HaloTag was induced with 20 µM IPTG for 2 hours and labeled with 40 pM JF549-HTL. Strains, *left*: control: bAB219, ↑*sepF*: bGS260, ↑*zapA*: bGS259. Strains, *right*: control: bAB309 and bGS104, ↑*sepF*: bGS158, ↑*zapA*: bGS159.

**Figure S4: Effects of EzrA overexpression on cell and Z ring morphology.**

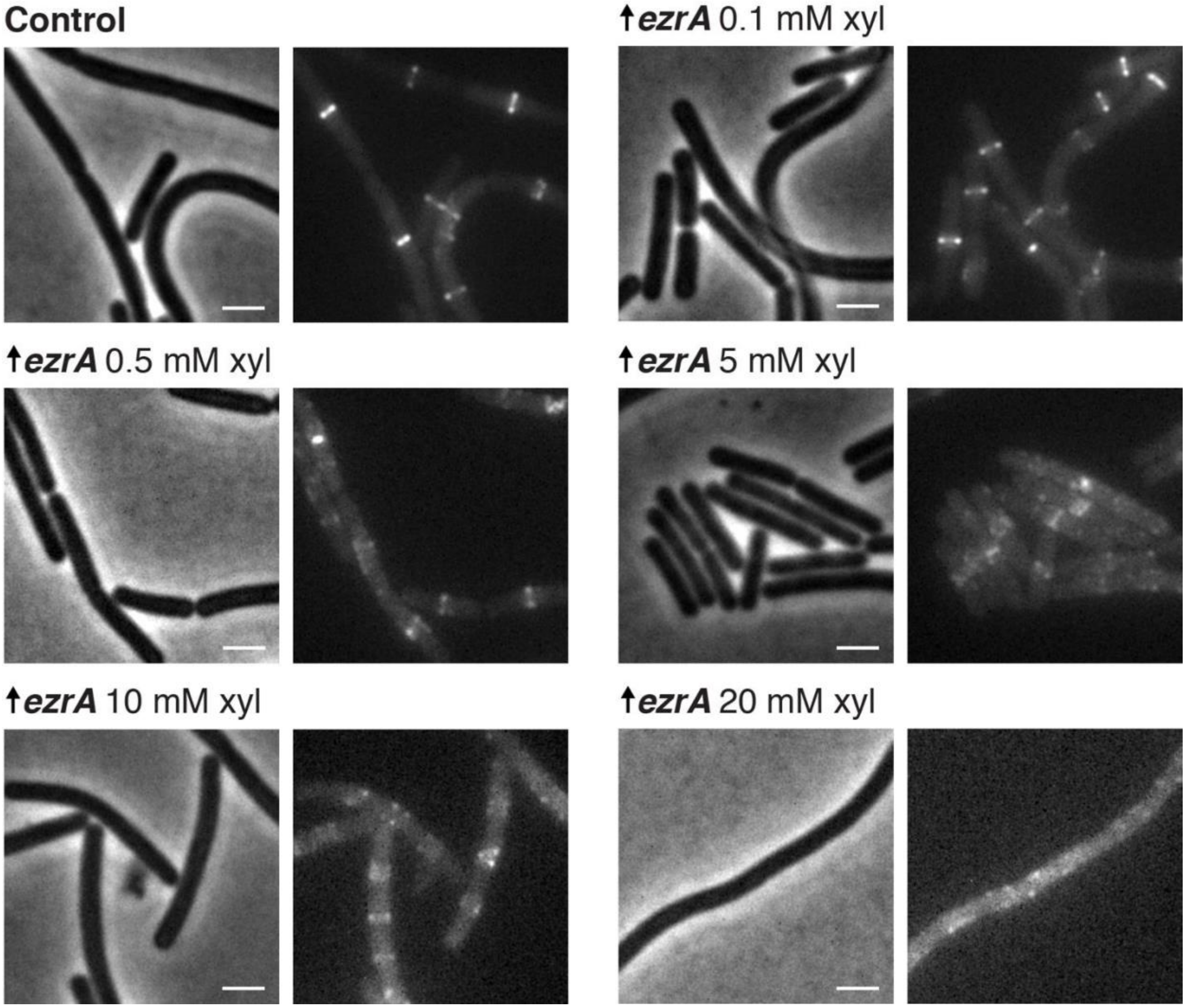


Each pair of images shows cell morphology (phase-contrast imaging, *left*), and Z ring morphology (epifluorescence images of cells expressing FtsZ-mNeonGreen induced with 20 µM IPTG for 2 hours, *right*), in control cells and cells with EzrA overexpressed. EzrA-overexpressing cells have perturbed Z ring morphology, as expected (*45*), a phenotype exacerbated with increasing induction. A second copy of *ezrA* was expressed from a xylose-inducible promoter by adding xylose at the indicated mM concentration. The 0.1, 0.5, and 5 mM concentrations were selected for quantitative analysis as the 10 and 20 mM xylose overexpression yielded unstable FtsZ filaments whose lifetimes were too short to be measured accurately. Strains: control: bAB219, *ezrA*↑: bGS263. Scale bars: 2 µm.

**Figure S5: Effects of removing synthetically lethal combinations of ZBPs on cell and Z ring morphology**

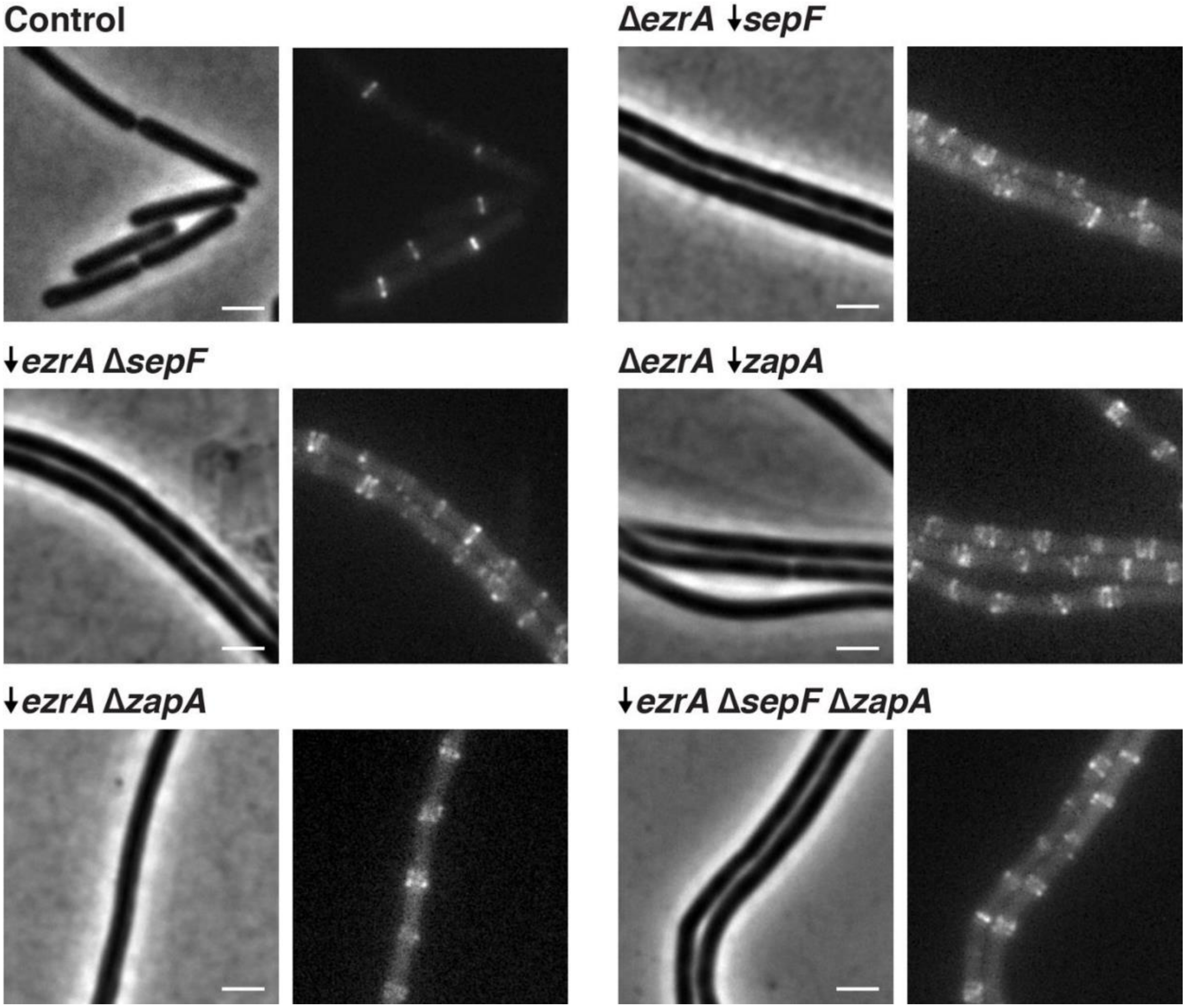


Each pair of images shows cell morphology (phase-contrast imaging, *left*), and Z ring morphology (epifluorescence images of cells expressing FtsZ-mNeonGreen induced with 20 µM IPTG for 2 hours, *right*), in control cells and cells lacking synthetically lethal combinations of ZBPs. To achieve this, a combination of knockouts (indicated by Δ) and depletions (indicated by ↓) were used; depletions were performed by expressing each gene under an inducible promoter until the start of the experiment, then withdrawing the inducer for 7 hours. This was repeated for all permutations of synthetically lethal combinations of ZBPs; all of these combinations result in elongated cells and disrupted Z ring architecture. Strains: control: bAB219, Δ*ezrA* ↓*sepF*: bGS290, ↓*ezrA* Δ*sepF*: bGS298, Δ*ezrA* ↓*zapA*: bGS293, ↓*ezrA* Δ*zapA*: bGS297, ↓*ezrA* Δ*sepF* Δ*zapA* (ΔZBPs): bGS308. Scale bars: 2 µm.

**Figure S6: Effects of removing synthetically lethal combinations of ZBPs on FtsZ**

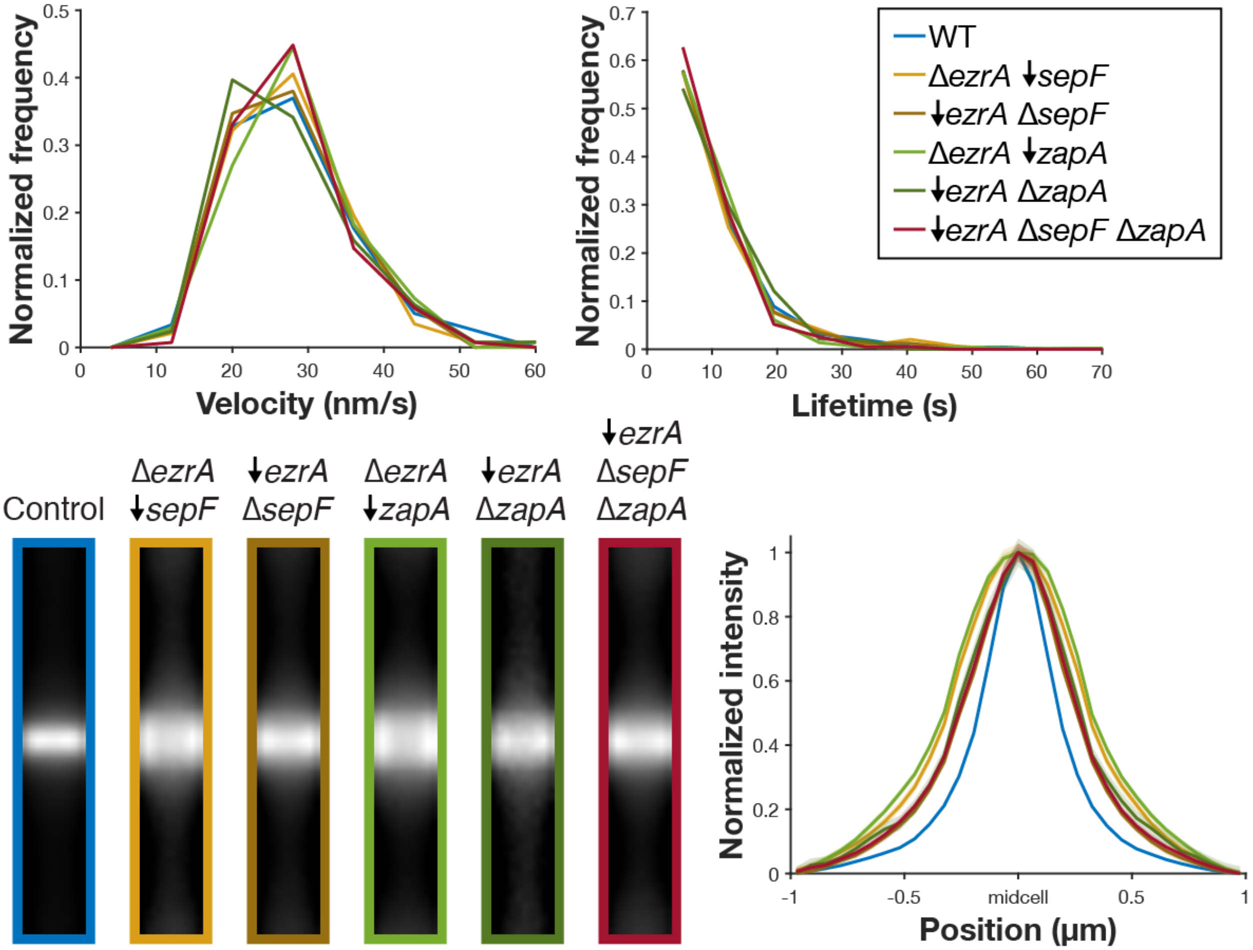


Velocity, lifetime, and Z ring morphology measurements for cells missing each synthetic lethal combination of ZBPs. All synthetic lethal combinations were investigated by a combination of knockouts (indicated by Δ) and depletions (indicated by ↓); depletions were performed by expressing the gene under an inducible promoter until the start of the experiment, then withdrawing the inducer for 7 hours.

**Top** Velocity (*left*) and lifetime (*right*) of cells missing synthetically lethal combinations of ZBPs are unchanged from control. For velocity measurements, FtsZ-mNeonGreen was induced with 20 μM IPTG for 2 hours, imaged by TIRFM, and then analyzed from kymographs. For lifetime measurements, FtsZ-HaloTag was induced with 20 μM IPTG for 2 hours and labeled with 40 pM JF549-HTL. Strains, *left*: control: bAB219, Δ*ezrA* ↓*sepF*: bGS290, ↓*ezrA* Δ*sepF*: bGS298, Δ*ezrA* ↓*zapA*: bGS293, ↓*ezrA* Δ*zapA*: bGS297, ↓*ezrA* Δ*sepF* Δ*zapA* (ΔZBPs): bGS308. Strains, *right*: control: bAB309 and bGS104, Δ*ezrA* ↓*sepF*: bGS204, ↓*ezrA* Δ*sepF*: bGS316, Δ*ezrA* ↓*zapA*: bGS206, ↓*ezrA* Δ*zapA*: bGS306, ↓*ezrA* Δ*sepF* Δ*zapA* (aka ΔZBPs): bGS331.

**Bottom** Z rings in cells missing synthetically lethal combinations of ZBPs are wider than control. Average intensity projections (*left*) and intensity profiles (*right*) of Z rings in each condition. Z rings were visualized using epifluorescence images of cells expressing FtsZ-mNeonGreen, induced with 20 μM IPTG for 2 hours. Z ring projections were created by averaging >100 Z ring images for each strain. Strains: control: bAB219, Δ*ezrA* ↓*sepF*: bGS290, ↓*ezrA* Δ*sepF*: bGS298, Δ*ezrA* ↓*zapA*: bGS293, ↓*ezrA* Δ*zapA*: bGS297, ↓*ezrA* Δ*sepF* Δ*zapA* (aka ΔZBPs): bGS308.

**Figure S7: Effects of a ∆*sepF* ∆*zapA* dual knockout**

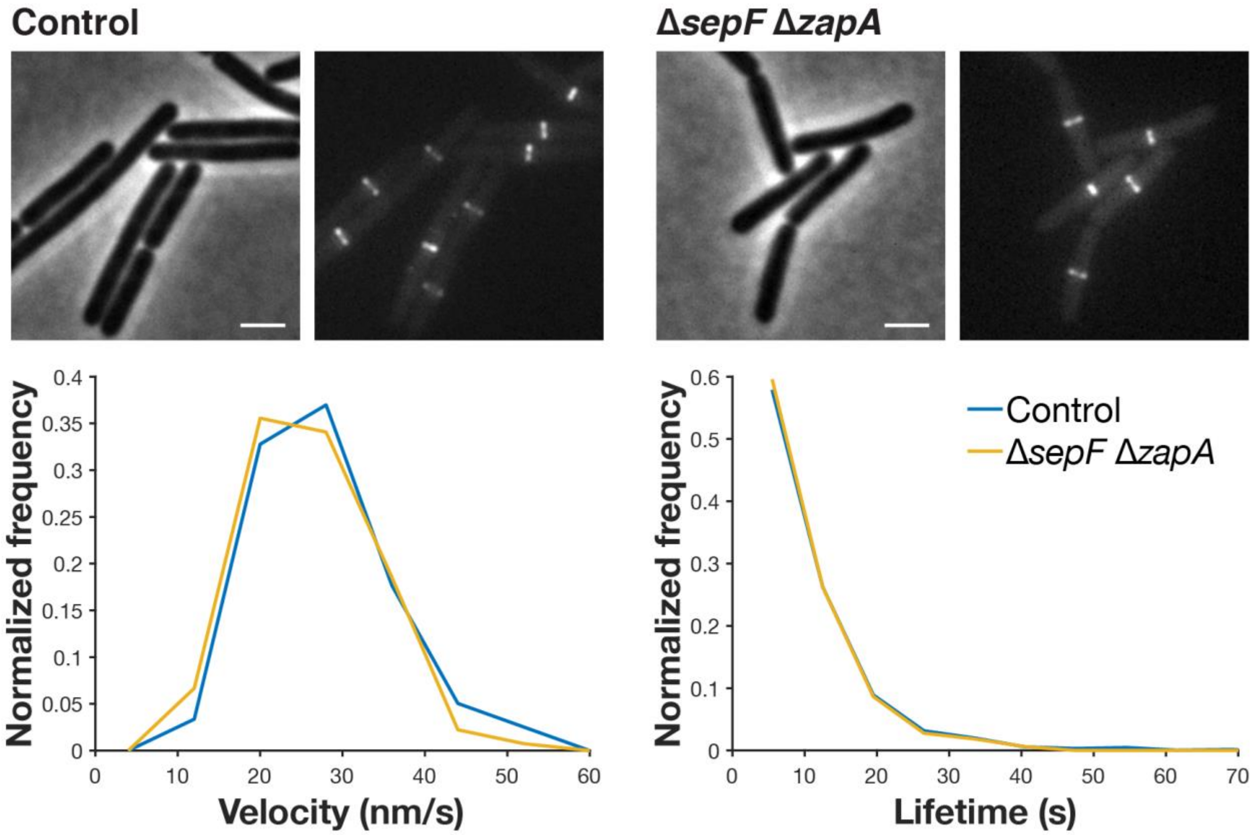


**Top** Each pair of images shows cell morphology (phase-contrast imaging, *left*), and Z ring morphology (epifluorescence images of cells expressing FtsZ-mNeonGreen induced with 20 µM IPTG for 2 hours, *right*), in control cells and cells with both *sepF* and *zapA* knocked out: ∆*sepF* ∆*zapA* is the only combination of ZBP deletions that is not synthetically lethal and their Z ring morphology is normal. Strains: control: bAB219, ∆*sepF* ∆*zapA*: bGS368. Scale bars: 2 µm.

**Bottom** ∆*sepF* ∆*zapA* cells have similar FtsZ treadmilling velocities (*left*) and subunit lifetimes (*right*) to control cells. For velocity measurements, FtsZ-mNeonGreen was induced with 20 µM IPTG for 2 hours, imaged by TIRFM, and analyzed from kymographs. For lifetime measurements, FtsZ-HaloTag was induced with 20 µM IPTG for 2 hours and labeled with 40 pM JF549-HTL. Strains, *left*: control: bAB219, ∆*sepF* ∆*zapA*: bGS368. Strains, *right*: control: bAB309 and bGS104, ∆*sepF* ∆*zapA*: bGS318.

**Figure S8: Characterization of the FtsZ(K86E) suppressor mutant**

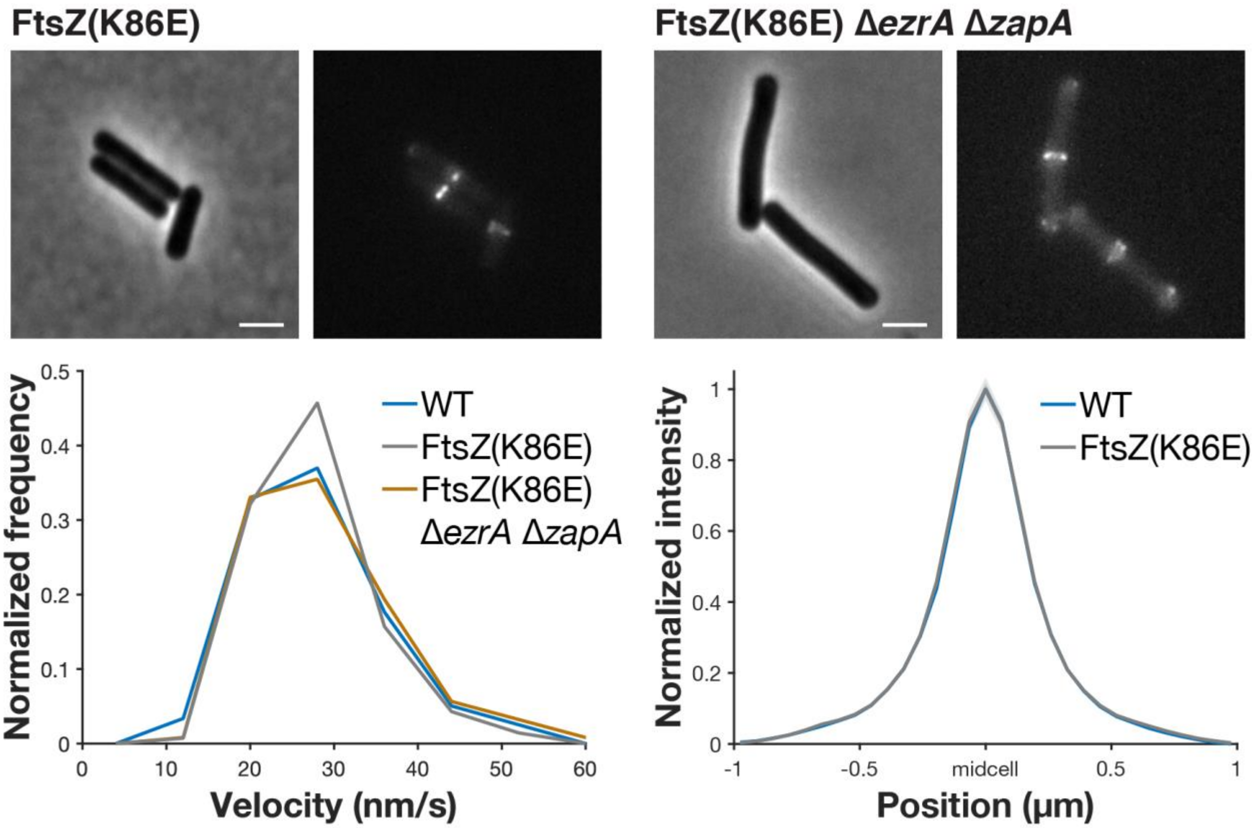


**Top** Each pair of images shows cell morphology (phase-contrast imaging, *left*), and Z ring morphology (epifluorescence images of cells expressing FtsZ-mNeonGreen induced with 20 µM IPTG for 2 hours, *right*) in FtsZ(K86E) and FtsZ(K86E) Δ*ezrA* Δ*zapA* cells. FtsZ(K86E) Z rings look similar to the control. Z rings in FtsZ(K86E) Δ*ezrA* Δ*zapA* are somewhat perturbed, but less so than typical cells missing synthetically lethal combinations of ZBPs; they also have polar Z rings, as is typical for Δ*ezrA* strains. Scale bars: 2 µm.

**Bottom** FtsZ(K86E) and FtsZ(K86E) Δ*ezrA* Δ*zapA* have similar FtsZ treadmilling velocities to control (*left*) and FtsZ(K86E) Z rings are identical in width to control, based on their intensity profiles (*right*). For velocity measurements in each strain, FtsZ-mNeonGreen was induced with 20 µM IPTG for 2 hours, imaged by TIRFM, and analyzed from kymographs. To generate Z ring intensity profiles, >100 Z ring images were averaged for each strain. Shaded area: SEM.

Strains: control: bAB219, FtsZ(K86E): bGS432, FtsZ(K86E) Δ*ezrA* Δ*zapA*: bGS463.

**Figure S9: Pbp2B localization and FDAA incorporation in ΔZBPs cells**

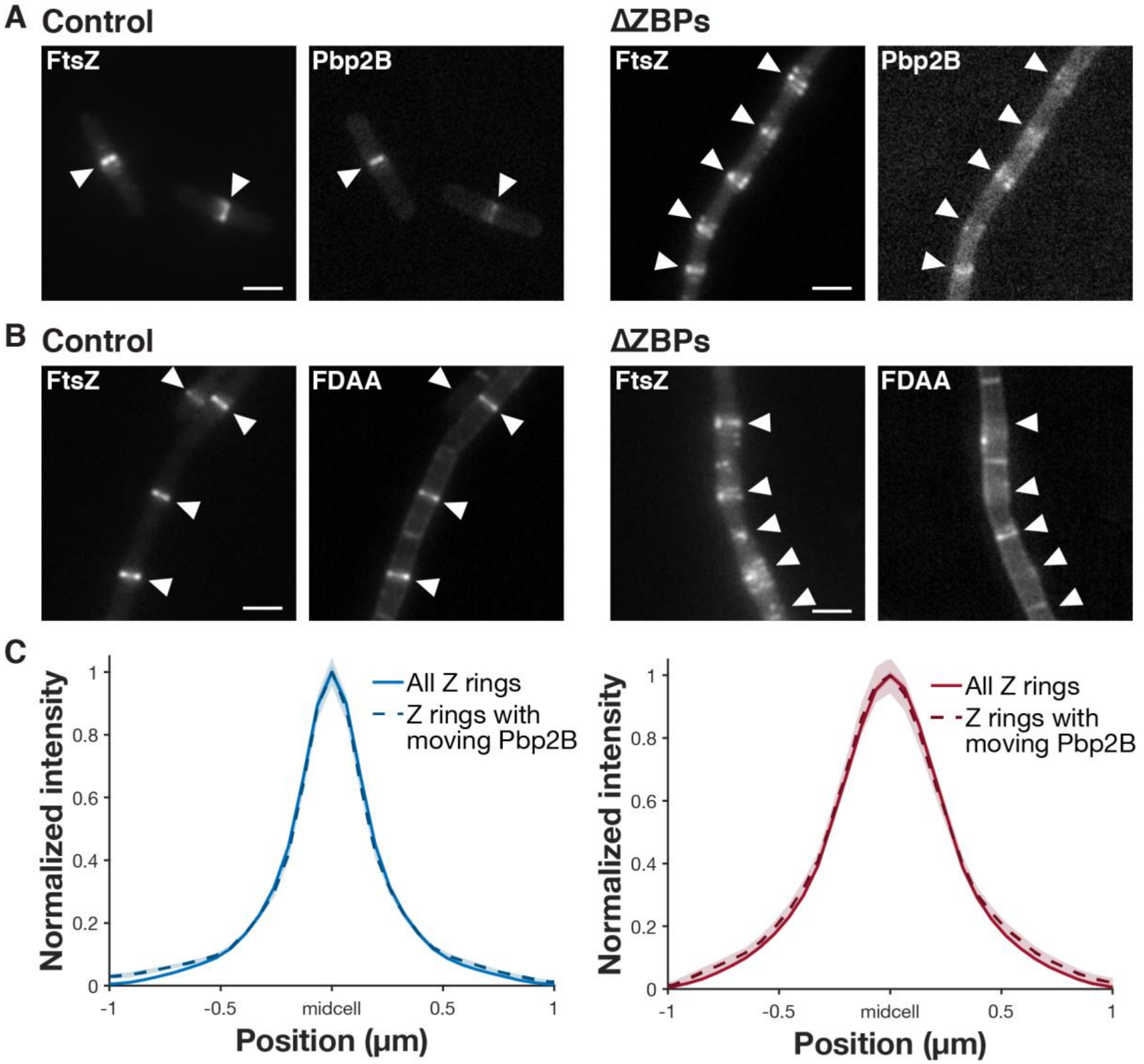


**A** Z rings (*left*) and Pbp2B localization (*right*) (epifluorescence images of cells expressing both Pbp2B-mNeonGreen and FtsZ-HaloTag induced with 20 µM IPTG for 2 hours and labeled with 5 nM JF549-HTL) in control and ΔZBPs cells. White arrows indicate the Z ring positions in each image. Strains: control: bGS104, ΔZBPs: bMH445. Scale bars: 2 µm.

**B** Z rings and fluorescent D-amino acid (FDAA) incorporation (epifluorescence images of cells labeled with 1 mM fluorescent D-lysine (FDL) for 30 seconds, *right*) in control and ΔZBPs cells. White arrows indicate the Z ring positions in each image. Strains: control: bMH510, ΔZBPs: bMH508. Scale bars: 2 µm.

**C** Pbp2B directional motion is seen at Z rings of all widths. The average intensity profiles of all Z rings (solid lines) and the Z rings at which Pbp2B moves directionally (dashed lines) for either control cells (*left*) or ΔZBPs cells (*right*) are similar. This confirms that the Pbp2B motion seen in ΔZBPs is present at decondensed rings. To generate intensity profiles of Z rings, >100 Z ring images were averaged for each strain. Shaded area: SEM. Strains: *left*: bMH512, *right*: bMH443.

### Table S1: Strains used in this study

| Description | Strain | Genotype | Source |
|---|---|---|---|
| *FtsZ single molecule imaging* | | | |
| FtsZ | bAB309 | *amyE::erm-Phyperspank-ftsA-HaloTag-15aa-ftsZ* | this study |
| 2 color | bGS104 | *amyE::erm-Phyperspank-ftsA-HaloTag-15aa-ftsZ, pbp2B::mNeonGreen-15aa-pbp2B* | this study |
| FtsZ(T111A) | bGS109 | *amyE::erm-Phyperspank-ftsA-HaloTag-15aa-ftsZ, ftsZΩftsZ(T111A) (tet)* | this study |
| MciZ expression | bGS328 | *amyE::erm-Phyperspank-ftsA-HaloTag-15aa-ftsZ, yvbJ::PxylA-mciZ::erm, pbp2B::mNeonGreen-15aa-pbp2B* | this study |
| Δ*zapA* | bGS141 | *amyE::erm-Phyperspank-ftsA-HaloTag-15aa-ftsZ, zapA-yshBD::tet* | this study |
| Δ*sepF* | bGS304 | *amyE::erm-Phyperspank-ftsA-HaloTag-15aa-ftsZ, sepF::scar* | this study |
| Δ*ezrA* | bGS167 | *amyE::erm-Phyperspank-ftsA-HaloTag-15aa-ftsZ, ezrA::cat* | this study |
| ↑*zapA* | bGS159 | *amyE::erm-Phyperspank-ftsA-HaloTag-15aa-ftsZ, ycgO::cat-pXyl-zapA* | this study |
| ↑*sepF* | bGS158 | *amyE::erm-Phyperspank-ftsA-HaloTag-15aa-ftsZ, ycgO::cat-pXyl-sepF* | this study |
| ↑*ezrA* | bGS157 | *amyE::erm-Phyperspank-ftsA-HaloTag-15aa-ftsZ, ycgO::cat-pXyl-ezrA* | this study |
| Δ*sepF* Δ*zapA* | bGS318 | *amyE::erm-Phyperspank-ftsA-HaloTag-15aa-ftsZ, zapA-yshBD::tet, sepF::scar* | this study |
| Δ*ezrA* ↓*sepF* | bGS204 | *amyE::erm-Phyperspank-ftsA-HaloTag-15aa-ftsZ, ezrA::scar, sepF::cat-pXyl-sepF* | this study |
| ↓*ezrA* Δ*sepF* | bGS316 | *amyE::erm-Phyperspank-ftsA-HaloTag-15aa-ftsZ, sepF::scar, ezrA::cat-pXyl-ezrA* | this study |
| Δ*ezrA* ↓*zapA* | bGS206 | *amyE::erm-Phyperspank-ftsA-HaloTag-15aa-ftsZ, ezrA::scar, zapA::cat-pXyl-zapA* | this study |
| ↓*ezrA* Δ*zapA* | bGS306 | *amyE::erm-Phyperspank-ftsA-HaloTag-15aa-ftsZ, zapA-yshBD::tet, ezrA::cat-pXyl-ezrA* | this study |
| ↓*ezrA* Δ*sepF* Δ*zapA* (ΔZBPs) | bGS331 | *amyE::erm-Phyperspank-ftsA-HaloTag-15aa-ftsZ, zapA-yshBD::tet, sepF::scar, ezrA::cat-pXyl-ezrA* | this study |
| *FtsZ filament and Z ring imaging* | | | |
| WT FtsZ | bAB219 | *amyE::erm-Phyperspank-ftsA-mNeonGreen-15aa-ftsZ* | (*5*) |
| FtsZ(T111A) | bAB281 | *amyE::erm-Phyperspank-ftsA-mNeonGreen-15aa-ftsZ, ftsZΩftsZ(T111A) (tet)* | (*5*) |
| MciZ expression | bGS326 | *amyE::erm-Phyperspank-ftsA-mNeonGreen-15aa-ftsZ, yvbJ::PxylA-mciZ::erm* | this study |
| Δ*zapA* | bGS250 | *amyE::erm-Phyperspank-ftsA-mNeonGreen-15aa-ftsZ, zapA-yshBD::tet* | this study |

| | | | |
|---|---|---|---|
| *ΔsepF* | bGS254 | *amyE::erm-Phyperspank-ftsA-mNeonGreen-15aa-ftsZ, sepF::tet* | this study |
| *ΔezrA* | bGS256 | *amyE::erm-Phyperspank-ftsA-mNeonGreen-15aa-ftsZ, ezrA::cat* | this study |
| *↑zapA* | bGS259 | *amyE::erm-Phyperspank-ftsA-mNeonGreen-15aa-ftsZ, ycgO::cat-pXyl-zapA* | this study |
| *↑sepF* | bGS260 | *amyE::erm-Phyperspank-ftsA-mNeonGreen-15aa-ftsZ, ycgO::cat-pXyl-sepF* | this study |
| *↑ezrA* | bGS263 | *amyE::erm-Phyperspank-ftsA-mNeonGreen-15aa-ftsZ, ycgO::cat-pXyl-ezrA* | this study |
| *ΔsepF ΔzapA* | bGS368 | *amyE::erm-Phyperspank-ftsA-mNeonGreen-15aa-ftsZ, sepF::scar, zapA-yshBD::tet* | this study |
| *ΔezrA ↓sepF* | bGS290 | *amyE::erm-Phyperspank-ftsA-mNeonGreen-15aa-ftsZ, ezrA::scar, sepF::cat-pXyl-sepF* | this study |
| *↓ezrA ΔsepF* | bGS298 | *amyE::erm-Phyperspank-ftsA-mNeonGreen-15aa-ftsZ, sepF::tet, ezrA::cat-pXyl-ezrA* | this study |
| *ΔezrA ↓zapA* | bGS293 | *amyE::erm-Phyperspank-ftsA-mNeonGreen-15aa-ftsZ, ezrA::scar, zapA::cat-pXyl-zapA* | this study |
| *↓ezrA ΔzapA* | bGS297 | *amyE::erm-Phyperspank-ftsA-mNeonGreen-15aa-ftsZ, zapA-yshBD::tet, ezrA::cat-pXyl-ezrA* | this study |
| *↓ezrA ΔsepF ΔzapA* (ΔZBPs) | bGS308 | *amyE::erm-Phyperspank-ftsA-mNeonGreen-15aa-ftsZ, sepF::scar, zapA-yshBD::tet, ezrA::cat-pXyl-ezrA* | this study |
| FtsZ(K86E) | bGS432 | *ftsZ::ftsZ(K86E), amyE::erm-Phyperspank-FtsA-mNeonGreen-15aa-FtsZ(K86E)* | this study |
| FtsZ(K86E) *ΔezrA ΔzapA* | bGS463 | *ftsZ::ftsZ(K86E), amyE::erm-Phyperspank-FtsA-mNeonGreen-15aa-FtsZ(K86E), ezrA::cat, zapA-yshBD::tet* | this study |
| *Other divisome proteins: single molecule imaging* | | | |
| EzrA | bMH42 | *ftsZ::mNeonGreen-15aa-ftsZ multicopy ezrA::ezrA-30aa-HaloTag-cat* | this study |
| SepF | bMH372 | *ftsZ::mNeonGreen-15aa-ftsZ multicopy amyE::erm-Phyperspank-sepF-30aa-HaloTag* | this study |
| ZapA | bMH49 | *ftsZ::mNeonGreen-15aa-ftsZ multicopy zapA::zapA-30aa-HaloTag-cat* | this study |
| DivIB | bAB366 | *ftsZ::mNeonGreen-15aa-ftsZ multicopy divIB::erm-Pxyl-HaloTag-15aa-divIB* | this study |
| DivIC | bAB367 | *ftsZ::mNeonGreen-15aa-ftsZ multicopy divIC::erm-Pxyl-HaloTag-15aa-divIC* | this study |
| FtsL | bGS165 | *ftsZ::mNeonGreen-15aa-ftsZ multicopy ftsL::erm-Phyperspank-HaloTag-15aa-ftsL* | this study |
| FtsW | bAB368 | *ftsZ::mNeonGreen-15aa-ftsZ multicopy ftsW::erm-Pxyl-HaloTag-15aa-ftsW* | this study |
| Pbp2B | bGS31 | *ftsZ::erm-mNeonGreen-15aa-ftsZ-cat multicopy pbp2b::erm-pHyperSpank-HaloTag-15aa-pbp2b* | (5) |
| *ZBP single molecule lifetimes* | | | |
| FtsA | bAB213 | *ftsAZ::erm-ftsA-HaloTag(sw)-ftsZ-cat multicopy* | (5) |
| EzrA | bMH03 | *ezrA::ezrA-30aa-HaloTag-cat* | this study |

| | | | |
|---|---|---|---|
| SepF | bMH332 | *amyE::erm-Phyperspank-sepF-30aa-HaloTag* | this study |
| ZapA | bMH28 | *zapA::zapA-30aa-HaloTag-cat* | this study |
| *Pbp2B dynamics* | | | |
| Control | bMH512 | *pbp2B::erm-Phyperspank-HaloTag-15aa-pbp2B, amyE::kan-Paz-ftsA-mNeonGreen-15aa-ftsZ* | this study |
| ↓*ezrA* Δ*sepF* Δ*zapA* (ΔZBPs) | bMH443 | *pbp2B::erm-Phyperspank-HaloTag-15aa-pbp2B, amyE::kan-Paz-ftsA-mNeonGreen-15aa-ftsZ, sepF::scar, zapA-yshBD::tet, ezrA::cat-pXyl-ezrA* | this study |
| *Pbp2B colocalization* | | | |
| Control | bGS104 | *see above* | this study |
| ↓*ezrA* Δ*sepF* Δ*zapA* (ΔZBPs) | bMH445 | *pbp2B::mNeonGreen-15aa-pbp2B, amyE::erm-Phyperspank-ftsA-HaloTag-15aa-ftsZ, sepF::scar, zapA-yshBD::tet, ezrA::cat-pXyl-ezrA* | this study |
| *Cell wall synthesis labeling* | | | |
| Control | bMH510 | *amyE::erm-Phyperspank-ftsA-HaloTag-15aa-ftsZ, dacA::kan* | this study |
| ↓*ezrA* Δ*sepF* Δ*zapA* (ΔZBPs) | bMH508 | *amyE::erm-Phyperspank-ftsA-HaloTag-15aa-ftsZ, sepF::scar, zapA-yshB::tet, ezrA::cat-pXyl-ezrA, dacA::kan* | this study |

Abbreviations: ↑: overexpression, ↓: depletion, 15aa: 15 amino acid linker, 30aa: 30 amino acid linker, (sw): sandwich fusion, scar: genomic scar remaining after an antibiotic cassette has been looped out, Paz: promoter of *ftsA-ftsZ* operon, multicopy: genotype of bAB185; see (*5*).

### Table S2: Constructs and plasmids used in this study

| ***Constructs from other studies*** | |
|---|---|
| **Construct** | **Reference** |
| *amyE::erm-Phyperspank-ftsA-mNeonGreen-15aa-ftsZ* | (*5*) |
| *ftsZ::mNeonGreen-15aa-ftsZ multicopy* | (*5*) |
| *ftsAZ::erm-ftsA-HaloTag(sw)-ftsZ-cat multicopy* | (*5*) |
| *pbp2B::mNeonGreen-15aa-pbp2B* | (*5*) |
| *pbp2b::erm-pHyperSpank-HaloTag-15aa-pbp2b* | (*5*) |
| *ftsZΩftsZ(T111A) (tet)* | (*5*) |
| *yvbJ::PxylA-mciZ::erm* | Gift from D. Z. Rudner |
| *zapA-yshBD::tet* | (*19*) |
| *dacA::kan* | (*51*) |

| ***Constructs created in this study*** | |
|---|---|
| **Amplicon** | **Primer: Sequence (5' to 3')** |
| *amyE::erm-Phyperspank-ftsA-HaloTag-15aa-ftsZ* | |
| amyE(up)-erm-Phyperspank-ftsA | oMD191: TTTGGATGGATTCAGCCCGATTG<br>oAB13: ccagtaccgatttctgccatGCTAAATCCTCCTAATCTGCCGAATG |
| HaloTag-15aa | oJE32: ATGGCAGAAATCGGTACTGG<br>oAB14: tggcctgagcccggtccctggccagatccctcgagGCCGCTGATTTCTAAGGTAGAAAG |
| 15aa-ftsZ-amyE(down) | oAB140: ggaccgggctcaggccaaggaagcggcATGTTGGAGTTCGAAACAAACATAGACG<br>oMD197: TCACATACTCGTTTCCAAACGGATC |
| *amyE::kan-Paz-ftsA-mNeonGreen-15aa-ftsZ* | |
| amyE(up)-kan | oMD191: TTTGGATGGATTCAGCCCGATTG<br>oSW42: TTCTGCTCCCTCGC |
| pAZ-ftsA(partial) | oAB76: gaacggtactgagcgagggagcagaaGTATTTGTTTCCGGTTTCT<br>oAB38: GCGAAGCTCTTCTGA |
| this assembly was transformed directly into bAB219 to complete the construct | |
| *ftsZ::ftsZ(K86E)-kan* | |
| ftsZ(up)-ftsZ(K86E) | oWM20: ATGAACAACAATGAACTTTACGTC<br>oWM66: cagggagcactggtcaactaccgttcgtatTTAGCCGCGTTTATTACGGT |
| kan | oSW40: CAGGGAGCACTGGTC<br>oSW42: TTCTGCTCCCTCGC |

| | |
|---|---|
| ftsZ(down) | oAB73: acattatacgaacggtactgagcgagggagcagaaTGTAAAGGACAAAATCGTTT<br>oAB30: CCATCCTCATATGTCTGACC |
| *amyE::erm-Phyperspank-FtsA-mNeonGreen-15aa-FtsZ(K86E)* | |
| amyE(up)-erm-Phyperspank | oMD191: TTTGGATGGATTCAGCCCGATTG<br>oMD232: GGTAGTTCCTCCTTAAAGCTTAATTGTTATCCGCTCACAAT |
| ftsA-mNeonGreen-15aa | oAB78: agcggataacaattaagctttaaggaggaactaccATGAACAACAATGAACTTTACGTC<br>oZB34: tggcctgagcccggtccctggccagatccctcgagcCTTATAGAGTTCATCCATACCCATC |
| 15aa-FtsZ(K86E) | oAB140: ggaccgggctcaggccaaggaagcggcATGTTGGAGTTCGAAACAAACATAGACG<br>oAB94: ctttcggtaagtcccgtctagccttgcccTTAGCCGCGTTTATTACGGTTTC |
| amyE(down) | oMD196: GGGCAAGGCTAGACGGG<br>oMD197: TCACATACTCGTTTCCAAACGGATC |
| *sepF::tet* | |
| sepF(up) | oMH43: TATTGGCCCGTCTATCAG<br>oMH98: gcgagggagcagaaCTCATTGCTGTACACCCC |
| tet | oSW40: CAGGGAGCACTGGTC<br>oSW42: TTCTGCTCCCTCGC |
| sepF(down) | oMH20: tgaccagtgctccctgAGCGAGATGATCCTTTATCAAG<br>oMH21: CTATGTATGAAGGATCTTCAACCA |
| *ezrA::cat* | |
| ezrA(up) | oMH53: GACATCTCCCGCTTGATG<br>oAB99: cgaacggtactgagcgagggagcagaaAATGAGCCCCCTTGCTGT |
| cat | oJM28: TTCTGCTCCCTCGCTCAG<br>oJM29: CAGGGAGCACTGGTCAAC |
| ezrA(down) | oMH05: tgaccagtgctccctgATAATCACGACCATGAAAAAGAG<br>oMH06: GTTGTGGATCGAGTCGGA |
| *ycgO::cat-pXyl-ezrA* | |
| ycgO(up) | oMD247: ATCGAACTGGCAAAAGGCAAAC<br>oMD248: tacgaacggtagttgaccagtgctccctgTCCCGCCATATAAATACAAATCGAAATAATC |
| cat-pXyl | oSW40: CAGGGAGCACTGGTC<br>oMD226: GGTAGTTCCTCCTTAATCGATCCATTCAAATACAGATGCATTTTATTTC |
| ezrA | oMH14: atcgattaaggaggaactaccATGGAGTTTGTCATTGGATTATTA<br>oGS37: acagccccttcctcctcctttcgatctCTAAGCGGATATGTCAGCTT |
| ycgO(down) | oMD257: AGATCGAAAGGAGGAGGAAGG<br>oMD252: CAAGGTTTTGAGCAGCTCAGTG |
| *ycgO::cat-pXyl-sepF* | |
| ycgO(up) | oMD247: ATCGAACTGGCAAAAGGCAAAC<br>oMD248: tacgaacggtagttgaccagtgctccctgTCCCGCCATATAAATACAAATCGAAATAATC |

| | |
|---|---|
| cat-pXyl | oSW40: CAGGGAGCACTGGTC<br>oMD226: GGTAGTTCCTCCTTAATCGATCCATTCAAATACAGATGCATTTTATTTC |
| sepF | oGS38: atggatcgattaaggaggaactaccATGAAAAATAAACTGAAAAACTTTTTCTCAATGG<br>oGS39: gggacagccccttcctcctcctttcgatctTTAGCCGCGTTTATTACGGTTTC |
| ycgO(down) | oMD257: AGATCGAAAGGAGGAGGAAGG<br>oMD252: CAAGGTTTTGAGCAGCTCAGTG |
| *ycgO::cat-pXyl-zapA* | |
| ycgO(up) | oMD247: ATCGAACTGGCAAAAGGCAAAC<br>oMD248: tacgaacggtagttgaccagtgctccctgTCCCGCCATATAAATACAAATCGAAATAATC |
| cat | oSW40: CAGGGAGCACTGGTC<br>oSW42: TTCTGCTCCCTCGC |
| pXyl-zapA | oSW38: cattatacgaacggtactgagcgagggagcagaaGAATTCGAGCTTGCATG<br>oGS36: acagccccttcctcctcctttcgatctTCAATCCTTTTCTTTAAGCTGACGC |
| ycgO(down) | oMD257: AGATCGAAAGGAGGAGGAAGG<br>oMD252: CAAGGTTTTGAGCAGCTCAGTG |
| *ezrA::cat-pXyl-ezrA* | |
| ezrA(up) | oMH35: GAATATGTCCGTCTCGCT<br>oMH54: tgaccagtgctccctgAATGAGCCCCCTTGCTG |
| cat-pXyl | oSW40: CAGGGAGCACTGGTC<br>oMD226: GGTAGTTCCTCCTTAATCGATCCATTCAAATACAGATGCATTTTATTTC |
| ezrA(partial) | oMH14: atcgattaaggaggaactaccATGGAGTTTGTCATTGGATTATTA<br>oMH56: CTTAGTACGGATTGACCGG |
| *sepF::cat-pXyl-sepF* | |
| sepF(up) | oAB109: GCCCGTGAGTATCACACG<br>oAB110: gctatacgaacggtagttgaccagtgctccctgACTCATTGCTGTACACCCCC |
| cat-pXyl | oSW40: CAGGGAGCACTGGTC<br>oMD226: GGTAGTTCCTCCTTAATCGATCCATTCAAATACAGATGCATTTTATTTC |
| sepF-<br>sepF(down) | oGS38: atggatcgattaaggaggaactaccATGAAAAATAAACTGAAAAACTTTTTCTCAATGG<br>oAB112: GCCAAAACCTCTGATAGACAGC |
| *zapA::cat-pXyl-zapA* | |
| zapA(up) | oMH22: AATGGCTTCAGGCTTTACTC<br>oMH58: tgaccagtgctccctgCGTTTCTCCTCCATTCCG |
| cat-pXyl | oSW40: CAGGGAGCACTGGTC<br>oMD226: GGTAGTTCCTCCTTAATCGATCCATTCAAATACAGATGCATTTTATTTC |
| zapA-<br>zapA(down) | oAB152: gtatttgaatggatcgattaaggaggaactaccTTGTCTGACGGCAAAAAAACA<br>oMH31: AGAGATTCTGCATCGTGT |
| *ezrA::ezrA-30aa-HaloTag-cat* | |
| ezrA(partial) | oMH01: GATTGCAAAGCTCAAGGATG<br>oMH02: AGCGGATATGTCAGCTTTGA |

| | |
|---|---|
| 30aa-HaloTag | oMH03: caaagctgacatatccgctCTTGAGGGTAGCGGACAAG<br>oMH04: agcgagggagcagaaTTAGCCGCTGATTTCTAAGGTAG |
| cat | oJM28: TTCTGCTCCCTCGCTCAG<br>oJM29: CAGGGAGCACTGGTCAAC |
| ezrA(down) | oMH05: tgaccagtgctccctgATAATCACGACCATGAAAAAGAG<br>oMH06: GTTGTGGATCGAGTCGGA |
| *amyE::erm-Phyperspank-sepF-30aa-HaloTag* | |
| amyE(up)-erm-pHyperSpank | oMD191: TTTGGATGGATTCAGCCCGATTG<br>oSW28: GGTAGTTCCTCCTTAAAGC |
| SepF-15aa-HaloTag | oMH45: ttaagctttaaggaggaactaccATGAGTATGAAAAATAAACTGAAAAACTT<br>oAB257: cggtaagtcccgtctagccttgcccTTAGCCGCTGATTTCTAAGG |
| amyE(down) | oMD196: GGGCAAGGCTAGACGGG<br>oMD197: TCACATACTCGTTTCCAAACGGATC |
| *zapA::zapA-30aa-HaloTag-cat* | |
| zapA(up) | oMH22: AATGGCTTCAGGCTTTACTC<br>oMH24: gtccgctaccctcaagATCCTTTTCTTTAAGCTGACGC |
| 30aa-HaloTag-cat | oMH25: CTTGAGGGTAGCGGACAA<br>oSW40: CAGGGAGCACTGGTC |
| zapA-zapA(down) | oMH29: tgaccagtgctccctgacaactATGCTAGATATCATCATC<br>oMH31: AGAGATTCTGCATCGTGT |
| *divIB::erm-Pxyl-HaloTag-15aa-divIB* | |
| divIB(up) | oAB235: GCCTGAGTATTTAAAGGCCATTG<br>oAB236: gtagttgaccagtgctccctgTGCCTGTTCACCTCATTCAA |
| erm-Pxyl-HaloTag-15aa | oJM29: CAGGGAGCACTGGTCAAC<br>oAB14: tggcctgagcccggtccctggccagatccctcgagGCCGCTGATTTCTAAGGTAGAAAG |
| 15aa-divIB-divIB(down) | oAB237: ctggccagggaccgggctcaggccaaggaagcggcATGAACCCGGGTCAAGAC<br>oAB238: CGCAAGCGATAAATAGTTTGAG |
| *divIC::erm-Pxyl-HaloTag-15aa-divIC* | |
| divIC(up) | oAB239: CGGCGTACACTAGCGAA<br>oAB240: gtagttgaccagtgctccctgACCAGACGGTCCTCCTTTC |
| erm-Pxyl-HaloTag-15aa | oJM29: CAGGGAGCACTGGTCAAC<br>oAB14: tggcctgagcccggtccctggccagatccctcgagGCCGCTGATTTCTAAGGTAGAAAG |
| 15aa-divIC-divIC(down) | oAB241: ctggccagggaccgggctcaggccaaggaagcggcTTGAATTTTTCCAGGGAACG<br>oAB242: CAGTGAATGCAAATGATGAGTC |
| *ftsL::erm-Phyperspank-HaloTag-15aa-ftsL* | |
| ftsL(up) | oMH49: CTTCTTCGTGAAACCGTAGA |

| | |
|---|---|
| | oMH50: tgaccagtgctccctgaGGCTGATGACCTCCTTTTA |
| erm-Phyperspank-HaloTag-15aa | oSW40: CAGGGAGCACTGGTC<br>oAB14: tggcctgagcccggtccctggccagatccctcgagGCCGCTGATTTCTAAGGTAGAAAG |
| 15aa-ftsL | oMH61: agggaccgggctcaggccaaggaagcggcATGAGCAATTTAGCTTACCAACC<br>oMH52: CGCTCCTTCAAATACTTATCCA |
| *ftsW::erm-Pxyl-HaloTag-15aa-ftsW* | |
| ftsW(up) | oME1: GAGAGACTTGATTATTTGCTTTCTTTTATC<br>oAB234: gtagttgaccagtgctccctgAACATCCTCTTCCCTGCTTC |
| erm-Pxyl-HaloTag-15aa | oJM29: CAGGGAGCACTGGTCAAC<br>oAB14: tggcctgagcccggtccctggccagatccctcgagGCCGCTGATTTCTAAGGTAGAAAG |
| 15aa-ftsW | oME6:ctcgagggatctggccagggaccgggctcaggccaaggaagcggcATGTTAAAAAAAATGCTAAAATCTTATGATTACTCAC<br>oME7: GTACACACTTGTTTTTTACAGATAAACAG |

Abbreviations: 15aa: 15 amino acid linker, 30aa: 30 amino acid linker, (sw): sandwich fusion, (up): homology region upstream of the indicated gene, (down): homology region downstream of the indicated gene, multicopy: genotype of bAB185; see (*5*). For primers, uppercase sequence indicates annealing region, and lowercase sequence indicates overhang.

**Table S3: FtsZ treadmilling velocity and velocities of other divisome proteins**

| Condition | Strain | Velocity (nm/s) | N |
|---|---|---|---|
| ***FtsZ treadmilling velocities*** | | | |
| Control | bAB219 | 28.5 ± 9.8 | 119 |
| Δ*ezrA* | bGS256 | 26.2 ± 8.1 | 132 |
| Δ*sepF* | bGS254 | 26.3 ± 7.5 | 120 |
| Δ*zapA* | bGS250 | 28.2 ± 6.9 | 125 |
| ↑*ezrA* 100 µM xyl | bGS263 + 100 µM xyl | 28.4 ± 8.5 | 134 |
| ↑*ezrA* 500 µM xyl | bGS263 + 500 µM xyl | 28.9 ± 9.9 | 128 |
| ↑*ezrA* 5 mM xyl | bGS263 + 5 mM xyl | 28.7 ± 9.3 | 128 |
| ↑*sepF* | bGS260 + 30 mM xyl | 27.7 ± 7.3 | 118 |
| ↑*zapA* | bGS259 + 30 mM xyl | 24.1 ± 6.6 | 110 |
| Δ*sepF* Δ*zapA* | bGS368 | 26.3 ± 7.3 | 132 |
| Δ*ezrA* ↓*sepF* | bGS290 | 28.2 ± 8.6 | 143 |
| ↓*ezrA* Δ*sepF* | bGS298 | 27.4 ± 7.6 | 121 |
| Δ*ezrA* ↓*zapA* | bGS293 | 28.2 ± 7.3 | 137 |
| ↓*ezrA* Δ*zapA* | bGS297 | 27.2 ± 8.0 | 126 |
| ↓*ezrA* Δ*sepF* Δ*zapA* (ΔZBPs) | bGS308 | 27.6 ± 7.1 | 136 |
| FtsZ(K86E) | bGS432 | 29.2 ± 9.2 | 123 |
| FtsZ(K86E) Δ*ezrA* Δ*zapA* | bGS463 | 27.8 ± 6.9 | 140 |
| ***Velocities of other divisome proteins*** | | | |
| DivIB | bAB366 | 26.2 ± 4.8 | 270 |
| DivIC | bAB367 | 26.7 ± 5.1 | 285 |
| FtsL | bGS165 | 25.4 ± 4.7 | 261 |
| FtsW | bAB368 | 24.1 ± 7.5 | 120 |
| Pbp2B | bGS31 | 25.6 ± 6.8 | 98 |

Velocity: mean ± standard deviation. N: number of trajectories analyzed. Abbreviations: ↑: overexpression, ↓: depletion, xyl: xylose.

### Table S4: Single-molecule lifetimes

| Condition | Strain | Lifetime (s) | N | p-value |
|---|---|---|---|---|
| ***FtsZ lifetimes*** | | | | |
| Control | bAB309, bGS104 | 8.1 (7.6, 8.7) | 1897 | |
| ↑*ftsAZ* | bAB309 + 100 µM IPTG | 8.8 (6.7, 13) | 455 | ns |
| FtsZ(T111A) | bGS109 | 16.5 (12.6, 23.8) | 337 | **** |
| MciZ expression | bGS328 | 3.9 (3.1, 5.1) | 442 | **** |
| Δ*ezrA* | bGS167 | 11.3 (7.8, 20.2) | 820 | **** |
| Δ*sepF* | bGS304 | 7.4 (6.5, 8.5) | 446 | ns |
| Δ*zapA* | bGS141 | 7.3 (6.4, 8.5) | 348 | ns |
| ↑*ezrA* 100 µM xyl | bGS157 + 100 µM xyl | 6.1 (5.5, 6.8) | 461 | *** |
| ↑*ezrA* 500 µM xyl | bGS157 + 500 µM xyl | 4.6 (4.2, 5) | 163 | **** |
| ↑*ezrA* 5 mM xyl | bGS157 + 5 mM xyl | 4 (3.6, 4.4) | 285 | **** |
| ↑*sepF* | bGS158 + 30 mM xyl | 6.1 (5.7, 6.7) | 338 | **** |
| ↑*zapA* | bGS159 + 30 mM xyl | 8.7 (7, 11.4) | 531 | * |
| Δ*sepF* Δ*zapA* | bGS318 | 7.8 (6.9, 8.9) | 324 | ns |
| Δ*ezrA* ↓*sepF* | bGS204 | 7.8 (6.7, 9.3) | 146 | ns |
| ↓*ezrA* Δ*sepF* | bGS316 | 8.1 (6.8, 10.2) | 399 | ns |
| Δ*ezrA* ↓*zapA* | bGS206 | 8.4 (5.8, 15.3) | 654 | ns |
| ↓*ezrA* Δ*zapA* | bGS306 | 9.4 (7.4, 13) | 281 | ns |
| ↓*ezrA* Δ*sepF* Δ*zapA* (ΔZBPs) | bGS331 | 7.4 (5.7, 10.3) | 193 | ns |
| 2 color: outside of Z ring | bGS104 | 5.6 (5.2, 6.1) | 383 | |
| 2 color: inside of Z ring | bGS104 | 11 (8, 17.4) | 238 | |
| 1 second intervals | bAB309 | 8.2 (7.3, 9.4) | 418 | ns |
| ***Lifetimes of other ZBPs*** | | | | |
| FtsA | bAB213 | 4.5 (3.9, 5.5) | 222 | **** |
| EzrA | bMH03 | 4.7 (4.1, 5.4) | 1160 | **** |
| SepF | bMH332 | 8 (6, 12) | 642 | *** |
| ZapA | bMH28 | 6.8 (5.6, 8.6) | 1056 | ns |

Lifetime: mean (95% confidence interval) from single exponential fit. N: number of particles analyzed. Abbreviations: ↑: overexpression, ↓: depletion, xyl: xylose. P-value computed from Wilcoxon rank-sum test vs control. ns $p \geq 0.05$, *$p < 0.05$, **$p < 0.01$, ***$p < 0.001$, ****$p < 0.0001$.

### Table S5: Z ring peak widths

| Condition | Strain | Z ring width (nm) | N |
| --- | --- | --- | --- |
| Control | bAB219 | 360 ± 40 | 2350 |
| Δ*ezrA* ↓*sepF* | bGS290 | 620 ± 60 | 638 |
| ↓*ezrA* Δ*sepF* | bGS298 | 510 ± 50 | 909 |
| Δ*ezrA* ↓*zapA* | bGS293 | 650 ± 60 | 1453 |
| ↓*ezrA* Δ*zapA* | bGS297 | 550 ± 50 | 183 |
| ↓*ezrA* Δ*sepF* Δ*zapA* (ΔZBPs) | bGS308 | 520 ± 50 | 812 |
| FtsZ(K86E) | bGS432 | 360 ± 40 | 840 |
| FtsZ(K86E) Δ*ezrA* Δ*zapA* | bGS463 | 510 ± 50 | 185 |

Z ring width: Full width at half maximum of the average Z ring intensity peak ± bootstrapped standard error. N: number of Z rings analyzed. Abbreviations: ↓: depletion.

### Supplemental Movies

**Movie S1:** FtsZ filaments treadmill around the cell. Cells expressing FtsZ-mNeonGreen (strain bAB219) were induced with 20 µM IPTG for 2 hours, then imaged at 1-second intervals for 100 seconds by TIRFM. The movie is displayed at 30 frames per second (30x actual speed). Scale bar: 5 µm.

**Movie S2:** Single molecules of EzrA, SepF, and ZapA are stationary. Each protein was expressed as a HaloTag fusion and labeled with JF549-HTL dye for single-molecule imaging using TIRFM. Movies begin with an image of the Z rings in each cell, visualized by FtsZ-mNeonGreen. Cells were imaged at 1-second intervals for 200 seconds; every other frame is displayed here. Strains: EzrA: bMH42, SepF: bMH372, ZapA: bMH49. The movie is displayed at 30 frames per second (60x actual speed). Scale bar: 5 µm.

**Movie S3:** Single molecules of DivIB, DivIC, FtsL, FtsW, and Pbp2B move directionally around the division site. Each protein was expressed as a HaloTag fusion and labeled with JF549-HTL dye for single-molecule imaging using TIRFM. Movies begin with an image of the Z rings in each cell, visualized by FtsZ-mNeonGreen. Cells were imaged at 1-second intervals for 200 seconds; every other frame is displayed here. Strains: DivIB: bAB366, DivIC: bAB367, FtsL: bGS65, FtsW: bAB368, Pbp2B: bGS31. The movie is displayed at 30 frames per second (60x actual speed). Scale bar: 5 µm.

**Movie S4:** Single-molecule imaging of FtsZ for lifetime analysis. Cells expressing FtsZ-HaloTag (strain bAB309) were imaged at 500-ms intervals for 50 seconds by TIRFM. The movie is displayed at 30 frames per second (15x actual speed). Scale bar: 5 µm.

**Movie S5:** EzrA overexpression decreases FtsZ filament length. The second copy of *ezrA* was expressed from a xylose-inducible promoter. Panel labels indicate the amount of xylose added in each experiment. Cells expressed FtsZ-mNeonGreen to visualized FtsZ filaments and were imaged at 1-second intervals for 100 seconds by SIM-TIRF microscopy. Strains: control: bAB219, ↑*ezrA*: bGS157. The movie is displayed at 30 frames per second (30x actual speed). Scale bar: 5 µm.

**Movie S6:** FtsZ filament treadmilling is unaltered in the absence of ZBPs. ΔZBPs cells have *sepF* and *zapA* knocked out, and *ezrA* depleted from a xylose-inducible promoter for 7 hours prior to imaging. Cells expressing FtsZ-mNeonGreen were imaged at 1-second intervals for 100 seconds by TIRF microscopy. Strains: control: bAB219, ΔZBPs: bGS308. The movie is displayed at 30 frames per second (30x actual speed). Scale bar: 5 µm.
